# Plasmin, the product of tissue plasminogen activator (tPA) treatment for ischemic stroke, impairs human brain endothelial barrier integrity

**DOI:** 10.64898/2026.05.27.728289

**Authors:** James Jack Willis Hucklesby, Cynthia Yaxin Gao, Euan Scott Graham, Catherine Elizabeth Angel

## Abstract

**Background:** tPA is used for the acute treatment of ischaemic stroke because it converts plasminogen to active plasmin, which breaks down clots. Previous studies show that tPA-activated plasminogen impairs brain endothelial barrier function. However, it is unclear whether the plasmin product of this reaction directly contributes to brain endothelial barrier deterioration.

**Objective:** Determine whether plasmin directly influences the human brain endothelial barrier.

**Methods:** We developed a new serum-free hCMEC/D3 culture model with ECIS real-time monitoring to establish how plasmin in isolation influences the brain endothelial barrier.

**Results:** ECIS monitoring demonstrated that plasmin caused a concentration-dependent decline in hCMEC/D3 barrier integrity, which was primarily mediated by a reduction in endothelial cell-to-cell interactions. Whilst a decrease in membrane capacitance and increase in basolateral adhesion were also observed, these changes were less marked. The inclusion of α2-antiplasmin ameliorated the changes in hCMEC/D3 barrier properties, suggesting this response is mediated by plasmin’s proteolytic activity. Quantitative immunocytochemistry confirmed that plasmin stimulated a decline in the key junctional molecules, Claudin-5, VE-Cadherin (CD144), β-Catenin, ZO-1 and PECAM-1 (CD31), which likely contributed to the deterioration of paracellular cell-to-cell interactions. Interestingly, using this serum-free model, tPA alone didn’t influence hCMEC/D3 barrier properties, whilst tPA with plasminogen did, implicating plasmin’s involvement.

**Conclusion:** Plasmin directly impaired the barrier function of hCMEC/D3 brain endothelial cell monolayers by stimulating a decline in key junctional molecules. This plasmin-mediated brain endothelial barrier deterioration has important implications for tPA use and should be considered whilst designing safer thrombolytic treatment options for patients experiencing acute ischemic stroke.

## INTRODUCTION

Acute ischemic stroke (AIS), caused by a blood clot in the cerebral microvasculature that restricts blood flow to the brain, remains a leading cause of death and disability worldwide [1,2]. Tissue-type plasminogen activator-based drugs (tPA) are the only FDA-approved thrombolytic agents for AIS with Alteplase, and more recently Tenecteplase, being the most widely used [3–5]. tPA converts the inactive precursor plasminogen into active plasmin. Plasmin then cleaves the fibrin strands scaffolding the blood clot to resolve the blockage [6]. It is recommended that tPA be limited to a therapeutic window of 4.5 hours following stroke onset, due to reduced efficacy outside this timeframe and increased risk of haemorrhagic transformation [2,7]. These limitations result in only a small proportion of patients receiving thrombolytic treatment [8]. Clinical trials for alternative pharmacological treatments have been largely unsuccessful, hence efforts to improve the safety and treatment window for current therapeutic options could be beneficial for patient outcomes.

The blood-brain barrier (BBB) is a key therapeutic target in AIS as BBB disruption is commonly observed in patients and is associated with haemorrhagic transformation [9]. While multiple cell types are crucial in supporting the BBB structure, this barrier is formed primarily by the endothelial cell layer lining the blood vessels, and the junctional molecules that bind them together [10,11]. There are two key classes of paracellular junctions: tight junctions (TJ) which regulate the permeability of the barrier, and adherens junctions which physically stabilize the barrier. Claudin-5 is the most prevalent TJ molecule at the BBB and is anchored to the cytoskeleton by Zono occludins [11]. Adherens junctions such as VE-cadherin anchor endothelial cells together, and link to the cytoskeleton via intracellular catenins e.g. β-catenin [11]. The cellular adhesion molecule PECAM-1 also contributes to BBB integrity, mechanosensing and leukocyte trafficking [12]. During ischemic stroke, these junctions are disrupted, and barrier permeability is increased, allowing bloodborne components to enter the brain and contribute to oedema and inflammation [9].

*In vivo* and clinical evidence links tPA treatment to endothelial disruption and an increased risk of haemorrhagic complications, however there are conflicting findings regarding the extent of damage mediated by plasmin, compared to signalling pathways activated directly by tPA [13–15]. A variety of plasmin independent effects of tPA have been proposed, including direct effects on neurons, astrocytes and pericytes, which subsequently influence endothelial barrier integrity. These mechanisms further complicate mechanistic interpretation [14,15], particularly given that plasmin may activate many of the same signalling pathways [13]. However, isolating plasmin’s direct effects remains challenging due to the ubiquitous presence of plasminogen system components *in vivo,* and in the serum used for most *in vitro* brain endothelial cell culture systems. Consequently, despite important therapeutic implications, the specific contribution of plasmin to BBB disruption remains unresolved [13].

To clarify the relative effects of plasmin and tPA on the brain endothelial barrier, we used the established hCMEC/D3 human brain microvascular endothelial cell line [16] with a novel serum-free cell culture medium. Electrical cell-impedance sensing (ECIS) technology was used to measure real-time barrier integrity, whereby tiny electrical currents are applied to the cell layer to measure it’s electrical resistance [17,18]. By measuring resistance at multiple frequencies, we were also able to mathematically model different brain endothelial barrier properties: R*_b_* represents the strength of endothelial cell-to-cell interactions, α represents the basolateral adhesion of endothelial cells to the underlaying substrate, and C*_m_* the endothelial cell membrane capacitance [17,18]. Collectively, this approach allowed us to introduce precise concentrations of individual plasminogen system components to the brain endothelial monolayer and monitor their effects on endothelial barrier properties in real-time. Quantitative immunocytochemistry (QICC) was then used to assess changes in key junctional proteins that mediate barrier function.

## METHODS

### Culture of Human Cerebral Microvascular Endothelial Cells (hCMEC/D3)

hCMEC/D3 cells were a kind gift from Pierre-Olivier Couraud [16]. All experiments used hCMEC/D3s between passages 30 and 40. hCMEC/D3s were cultured in flasks (Falcon, 353136) coated with 150 µg/ml collagen I (Invitrogen, C3867). hCMEC/D3s were cultured in EBM-2MV serum containing media (5% FCS, PromoCell, C-22121) at 37°C, with 5% CO_2_ and 100% humidity, as previously described [16,19]. Cells at ∼90% confluence were passaged using TrypLE (Life Technologies, 12604021).

Cell suspensions were cryopreserved in 90% FBS (Moorgate) and 10% DMSO (Sigma-Aldrich, D2650). Aliquots in cryovials (Nunc, 377267) were cooled to −80°C (CoolCellLX, BioCision, BCS-405) and transferred to vapour phase liquid nitrogen. Cells were thawed at 37°C, resuspended in EBM-2MV serum containing media and transferred to a collagen-coated flask for culture. Cultures were routinely tested for mycoplasma (LiliF Diagnostics, 25237).

#### Preparation of AIM-V serum-free media for *hCMEC/D3s*

AIM-V serum-free media was prepared by supplementing AIM-V media (ThermoFisher Scientific, 12055091) with 1X GlutaMAX (ThermoFisher Scientific, 35050061) and 2MV supplements (PromoCell, C-39221 with FCS omitted) and stored at −80°C. Before use, media was thawed at 37°C; remaining media was stored at 4°C and used within 7 days.

#### Preparation of plasminogen system components

Plasmin (HPLM-1.0MG), plasminogen (HGPG-1.0MG) and α2-antiplasmin (HA2AP-1.0MG) proteins purified from human plasma were purchased in lyophilised form from Molecular Innovations. tPA (Actilyse) was kindly donated by Boehringer Ingelheim Ltd (NZ). Proteins were reconstituted to pre-lyophilisation volumes in sterile Type I water, gently re-suspended, aliquoted into low-bind tubes (Eppendorf, EP0030108116), snap-frozen in liquid nitrogen and stored at −80°C. After thawing, these protein treatments were prepared in low-bind tubes and used immediately.

#### D-VLC-AMC plasmin fluorescent substrate assay

The plasmin-specific substrate H-D-Val-Leu-Lys-AMC (Bachem, 4008009, henceforth referred to as D-VLC-AMC) was reconstituted in DMSO and stored at −80°C in single-use aliquots. 10X substrate solution was prepared in DMEM (Life Technologies, 10313021) and 10µl was pipetted into each well of a clear bottom black-sided 96-well plate (Corning, 3603). Next, 10uL of plasminogen system component in DMEM was added to the wells, followed by 80uL of test media, or DMEM if not specified. The plate was sealed (PerkinElmer, 6050185) and incubated at 37°C in a SpectraMax iD3 plate reader (with 10 second stir at start); fluorescence readings were taken at an excitation of 380 nm and emission of 460 nm every 5 minutes for 16 hours. Data were analysed using RStudio, and graphed with ggplot2 [20]. Data were presented as percentages of the 100nM plasmin positive control measured at 16hr.

#### Measuring brain endothelial barrier properties in real-time using Electric Cell-Substrate Impedance Sensing (ECIS)

ECIS 96W20idf plates (Applied BioPhysics) were coated with 10mM L-cysteine (Sigma-Aldrich, C7352). Wells were subsequently coated with 150µg/ml collagen I. Brain endothelial cells were seeded at a density of 62,500 cells/cm^2^ in 150µl EBM-2MV in each well. The plate was then placed in the ECIS 96 well array station, housed in a cell culture incubator at 37°C, 5% CO_2_. ECIS captured impedance data using the Multiple Frequency Time-series Mode for the duration of the experiment.

To apply treatments, ECIS monitoring was paused, the plate removed, treatments were applied in a cell culture hood. Plates were then reinstalled into the ECIS array station and monitoring resumed.

The multifrequency ECIS impendence data was modelled using ECIS Software (Applied Biophysics, V 1.1.252). This modelled impedance data represents three different brain endothelial barrier properties: R*_b_* represents the electrical resistance of the paracellular component of the barrier i.e. the cell-to-cell junctions; alpha represents the resistance between the basal cell membrane and the electrode on the bottom of the well i.e. basolateral adhesion of the cells to the collagen substrate that mimics the basement membrane; C*_m_* represents the capacitance of the cell membrane [21]. ECIS data was imported into vascr (V 0.1.6), which statistically assessed and graphed data [22].

#### Quantifying brain endothelial junctional molecule expression using immunocytochemistry with vjunctr

hCMEC/D3 cells were seeded into Ibidi µ-slide 96 well plates (Ibidi, 89626) at 62,500 cells/cm^2^ in 200uL EBM-2MV per well, cultured for 48 hours, washed and transferred to AIM-V serum-free media (if not otherwise specified) for 16 hours. Treatments were then applied. Following culture, cells were fixed with 4% formaldehyde in PBS at room temperature for 10 minutes, washed once with PBST (1X PBS Tablet (ThermoFisher, 18912014) in 500ml deionised water + 0.1% Tween 20 (Sigma-Aldrich, 93773)) and again with 0.1% Triton X-100 (Sigma-Aldrich, X100) in PBS for 10min. Cells were washed thrice with PBST, then blocked with ICC blocking buffer (PBST with 1% BSA (pH Scientific, PH700-100g)) for 10 minutes at room temperature. Cells were then probed with primary antibodies (VE-Cadherin clone F8, Santa Cruz; β-Catenin clone 5H10, Invitrogen; Claudin-5 clone EPR7583, AbCam; ZO-1 clone ZO1-1A12, Invitrogen; PECAM-1 clone WM59, BioLegend) in ICC blocking buffer for one hour at room temperature. Thereafter, cells were washed thrice for 5 minutes each with PBST. Cells were then incubated with the respective fluorophore conjugated secondary antibodies (goat anti-rabbit Alexa-568 (ThermoFisher, A-11011) or anti-mouse Alexa-568 (ThermoFisher, A21124, 1:200)) & DAPI (Invitrogen, D1306) in ICC blocking buffer at room temperature for one hour. Cells were washed thrice with PBST, transferred to ICC storage buffer (PBS + 0.05% Sodium Azide (Merck, S2002). Images were acquired using an Operetta CLS High-Content Imaging System (PerkinElmer), with 20x/ 0.75 NA lens. Single plane images from nine fields per well were captured. The total and junctional florescence intensity was statistically assessed using vjunctr (Sup Fig 5) and plotted using ggplot2 [20].

## RESULTS

Initially we conducted fluorogenic substrate assays, which confirmed that the commercial plasmin is enzymatically active, is effectively inhibited by α2-antiplasmin, that plasminogen is inactive alone, and that tPA efficiently converts plasminogen into active plasmin [23] (Sup. Fig. 1).

### The widely used EBM-2MV serum-containing media is not suitable to study the direct effect of plasmin on hCMEC/D3 cells, however AIM-V-2MV serum-free media is suitable

To investigate the direct effects of plasmin on brain endothelial cells, we first needed to ensure that the media being used did not contain plasmin, or any plasminogen system components which influence plasmin activity. To identify plasmin activity, we again used the D-VLC-AMC plasmin-specific fluorogenic substrate assay. First, we assessed the standard EBM-2MV serum-containing media widely used for hCMEC/D3 culture (Fig 1A & D). EBM-2MV serum-containing media alone generated a detectable fluorescent signal, indicating the presence of endogenous plasmin activity. Additionally, the addition of exogenous tPA generated a detectable fluorescent signal, indicating the presence of plasminogen within the EBM-2MV serum-containing media [6]. Finally, the enzymic activity of exogenous plasmin in EBM-2MV serum-containing media was substantially lower than that detected in the serum-free counterpart (Fig 1A & B, D & E), suggesting the presence of plasmin inhibitors in the serum-containing formulation. Collectively these data demonstrate that EBM-2MV serum-containing media, contains endogenous plasmin, plasminogen, and plasmin inhibitors and hence does provide the plasminogen system free conditions required to study the direct effect of exogenous plasmin on hCMEC/D3 brain endothelial cells.

**Figure 1.**
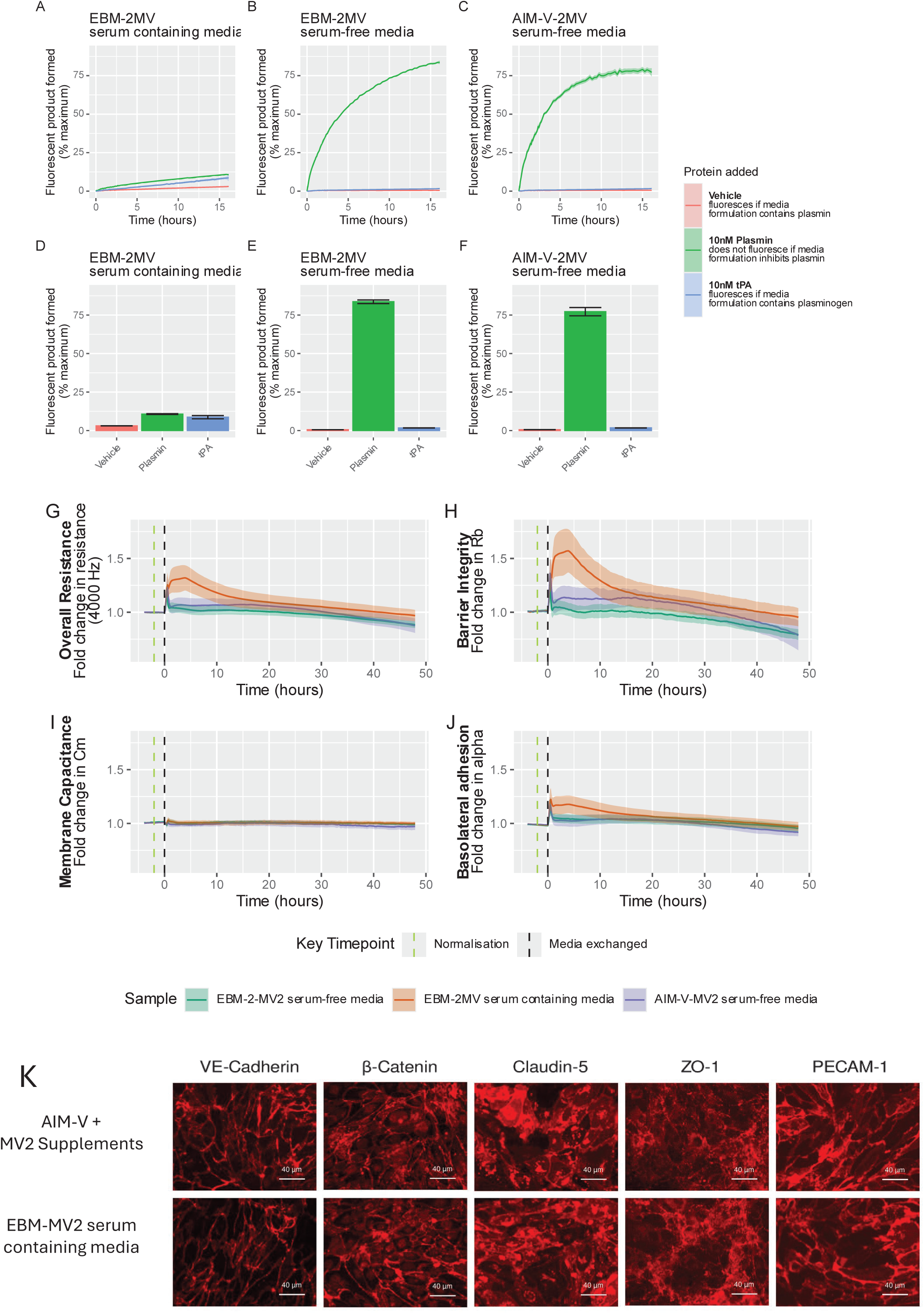
AIM-V-2MV serum-free media does not contain plasminogen system components or interact with it and can maintain a stable functional hCMEC/D3 brain endothelial barrier. Initially D-VLC-AMC plasmin fluorescent substrate assays were conducted to identify a culture media that does not contain plasminogen system components, so that the effect of plasmin on hCMEC/D3 could be assessed in isolation. The standard EBM-2MV serum containing media for hCMEC/D3, contains plasmin, plasminogen and factors that inhibit plasmin activity, and hence cannot be used for this study (A). serum-free EBM and EBM + 2MV supplements do not contain plasmin, plasminogen or factors that inhibit plasmin activity (A); however, they do contain an insufficient level of protein (Sup Fig 2), which is not physiologically representative and furthermore is unlikely to support low quantities of plasmin protein in solution. The base media was therefore changed to AIM-V, which contains a higher level of protein. AIM-V + 2MV supplements does not contain plasmin or plasminogen, and does not inhibit plasmin activity, and hence is suitable for studying the direct effect of exogenous plasmin on hCMEC/D3 (C). The data presented are the mean ± SEM of three independent experiments. ECIS monitoring demonstrates that AIM-V-2MV serum-free media supports the maintenance of a stable functional hCMEC/D3 brain endothelial barrier, with similar properties to hCMEC/D3 layers cultured in EBM-2MV serum-free media (B, C, E, & F). Briefly hCMEC/D3 cells were seeded into ECIS 96W20IDF plates and cultured in EBM-2MV serum containing media for 48 hours to allow a stable barrier to form. Cells were then washed thrice with their respective future culture media (i.e. AIM-V-2MV serum-free media, EBM-2MV serum-free media or EBM-2MV serum containing media). Cells were then cultured in their respective media for a further 40 hours. Barrier properties were monitored in real-time throughout the experiment using ECIS. Ribbon plots represent the mean ± SEM of data from three independent experiments, each consisting of three technical replicates. Data was normalised before treatments were applied using vascr. Immunocytochemistry illustrates that hCMEC/D3 brain endothelial cells cultured in AIM-V-2MV serum-free media express the key junction associated molecules VE-cadherin (CD144), *p*-Catenin, Claudin-5, ZO-1 and PECAM-1 (CD31) (D). Briefly hCMEC/D3 cells were cultured in Ibidi 96 well plates in EBM-2MV serum containing media for 48 hours. Cells were then washed thrice with their respective future culture media (i.e. AIM-V-2MV serum-free media or EBM-2MV serum containing media) and cultured in their respective media for 16 hours. Cells were fixed with paraformaldehyde, probed with antibodies detecting junctional molecules, washed, and probed with respective fluorophore conjugated secondary antibodies and DAPI. Cells were imaged using an Operetta CLS. Images are representative of 9 fields from each of three independent experiments.

In contrast, EBM-2MV serum-free media showed no detectable plasmin activity and little increase in fluorescence following tPA addition (Fig. 1B & E), indicating a negligible level of endogenous plasmin and plasminogen respectively. Exogenous plasmin added to EBM-2MV serum-free media caused robust substrate cleavage, showing that this media formulation does not or minimally inhibits plasmin activity (Fig. 1B & E). Collectively therefore, EBM-2MV serum-free media contains no or very little endogenous plasmin, plasminogen, or plasmin inhibitors. Despite this, the EBM-2MV serum-free media formulation is not suitable for subsequent studies as it contains insufficient total protein to maintain low concentrations of exogenously added plasminogen system components in solution (Sup. Fig. 2) [24].

We therefore evaluated the viability of using the 2MV-supplement with AIM-V media, which contains a high level of defined proteins, whilst remaining serum-free (Fig. 1E & F). AIM-V-2MV serum-free media exhibited no detectable plasmin activity, which did not increase with tPA’s inclusion, confirming very low levels of plasmin and plasminogen respectively. Furthermore, addition of exogenous plasmin caused robust D-VLC-AMC cleavage, indicating that AIM-V-2MV inhibition of plasmin activity is negligible. Together, these data demonstrate that AIM-V-2MV serum-free media lacks plasminogen system components and does not interfere with plasmin activity, making it suitable for studying the direct effect of exogenous plasmin on cells.

### AIM-V-2MV serum-free media supports a functional hCMEC/D3 endothelial barrier stabilised by key junctional molecules

We next assessed whether the AIM-V-2MV serum-free media could maintain a functional brain endothelial barrier (Fig. 1G-J). Real-time ECIS revealed that hCMEC/D3 monolayers transferred from EBM-2MV serum containing media into AIM-V-2MV serum-free media maintained a stable functional barrier for 48 hrs. Resistance, cell-to-cell interactions (R*_b_*), membrane capacitance (C*_m_*), and basolateral adhesion (alpha) were comparable to those observed for hCMEC/D3 monolayers maintained in EBM-2MV serum-free media (Fig. 1G-J). Furthermore, hCMEC/D3 monolayers transferred into serum-free AIM-2MV did not experience the immediate temporal increase in resistance, R*_b_* and alpha, that is routinely observed when brain endothelial cells are provided with fresh serum containing media [17,18].

Immunocytochemical analysis confirmed that hCMEC/D3 cells transitioned into AIM-V-2MV serum-free media retained key endothelial junction-associated proteins, including VE-cadherin, β-catenin, Claudin-5, ZO-1, and PECAM-1 (Fig. 1K). Localisation of these junctions mirrored hCMEC/D3s cultured in EBM-2MV serum-containing media and aligned with the preserved paracellular barrier properties (R*_b_*) detected by ECIS (Fig. 1K).

Given plasmin’s enzymatic activity, we next assessed whether plasmin could cleave components of AIM-V-2MV serum-free media and impair its ability to support hCMEC/D3 barrier function (Sup. Fig. 3). Despite potential susceptibility predicted from sequence information *in silico*, plasmin pre-treated media retained its capacity to support a stable hCMEC/D3 barrier (Sup. Fig. 3B). Furthermore, to ensure responses in future experiments were attributable solely to exogenous plasmin treatment, we used fluorogenic assays to confirm that live hCMEC/D3 cells transitioned to AIM-V-2MV media retained minimal cell surface-bound plasminogen (Sup. Fig. 4).

Collectively, these data demonstrate that AIM-V-2MV serum-free media can support a stable and functional hCMEC/D3 brain endothelial barrier and there is minimal contaminating plasmin or plasminogen in this culture model. Hence, this represents a suitable model to study the direct effect of plasmin on human brain endothelium.

### tPA alone does not influence barrier properties, whilst tPA activated plasminogen does impair barrier properties of hCMEC/D3 brain endothelium in AIM-V-2MV serum-free media

Next, we investigated whether therapeutically relevant circulating concentrations of tPA (10 nM) [25], in the presence and absence of plasminogen, influenced brain endothelial barrier integrity in our model (Fig. 2). Across all ECIS parameters, including resistance, R*_b_*, C*_m_*, and alpha, no significant differences were observed between the hCMEC/D3 layers treated with vehicle and tPA (Fig. 2A-L); both tPA and vehicle exhibited comparable transient responses before barrier properties stabilised and remained closely aligned for the duration of the experiment. In contrast, simultaneous addition of tPA and plasminogen caused a significant sustained decline in overall resistance and cell-to-cell interactions (R*_b_*) (Fig. 2A-F), and a lesser decline in basolateral adhesion (alpha) relative to the vehicle (Fig. 2J-L). Interestingly the tPA with plasminogen response was multiphasic. Following the initial decrease, the resistance and R*_b_* plateaued below the vehicle for ∼4 hours, thereafter there was a further sustained reduction in resistance and R*_b_* relative to the vehicle for the remainder of the experiment (Fig. 2A & D). tPA and plasminogen stimulated C*_m_* to initially increase above the vehicle, plateau for ∼6 hours, then realign and slowly elevate with the vehicle (Fig. 2G).

**Figure 2.**
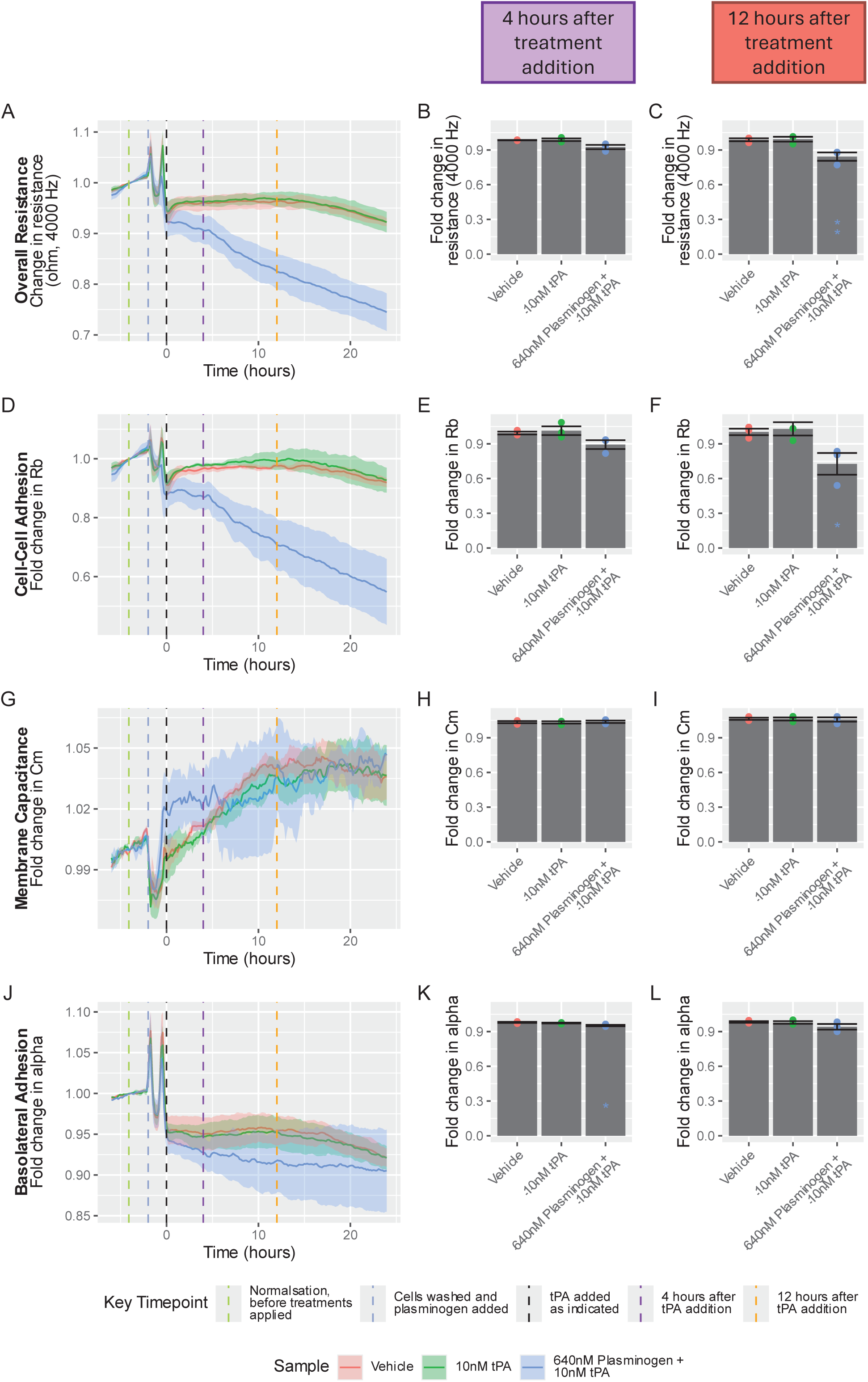
tPA in AIM-V-2MV serum-free media does not influence hCMEC/D3 brain endothelial barrier properties, whilst tPA with plasminogen does impair barrier integrity. hCMEC/D3 cells were seeded into ECIS 96W20IDF plates and cultured in EBM-2MV serum containing media for 48 hours to allow a stable barrier to form. Cells were then washed thrice with AIM-V-2MV serum-free media and cultured for 16 hours in AIM-V-2MV serum-free media. Cells were washed thrice and cultured for 1 hour in fresh AIM-V-2MV serum-free media ± 640nM plasminogen. tPA was then added to the respective wells and cells cultured for a further 24 hours. Barrier properties were monitored in real-time throughout the experiment using ECIS. Ribbon plots represent the mean ± SEM of data from three independent experiments, each consisting of three technical replicates. Data was normalised 1hr before plasminogen was added, using vascr. Statistical analysis of this data was carried out at 4 and 12 hours after tPA addition; each treatment response was compared with vehicle control using two-way ANOVA with Dunnett’s test (* p <0.05, ** p <0.01).

These data demonstrate that in the absence of plasminogen, tPA alone does not impact hCMEC/D3 brain endothelial barrier properties when cells are cultured in AIM-V-2MV serum-free media. However, activation of plasminogen with tPA causes significant barrier deterioration, predominantly mediated by disruption of the cell-to cell junctions.

### Plasmin disrupts the hCMEC/D3 brain endothelial barrier in serum-free media

To determine the direct impact of plasmin on brain endothelial barrier integrity, hCMEC/D3 monolayers cultured in AIM-V-2MV serum-free media were treated with increasing concentrations of plasmin, and barrier properties were monitored in real time using ECIS (Fig. 3). Following plasmin addition, a progressive and concentration-dependent reduction in overall barrier resistance was observed during the 24-hour treatment period, with higher plasmin concentrations (≥160nM) causing low level but statistically significant barrier deterioration at 4 hours, and 640nM remaining significant at 12 hours (Fig. 3A-C). The response to ≥320nM plasmin was multiphasic; following the initial treatment peak, there was a rapid decline during the first hour, resistance plateaued for ∼2-3 hours and then steadily declined until the end of the experiment (Fig. 3A). Lower plasmin concentrations (≤80nM) produced no significant change in barrier resistance compared with vehicle controls.

**Figure 3.**
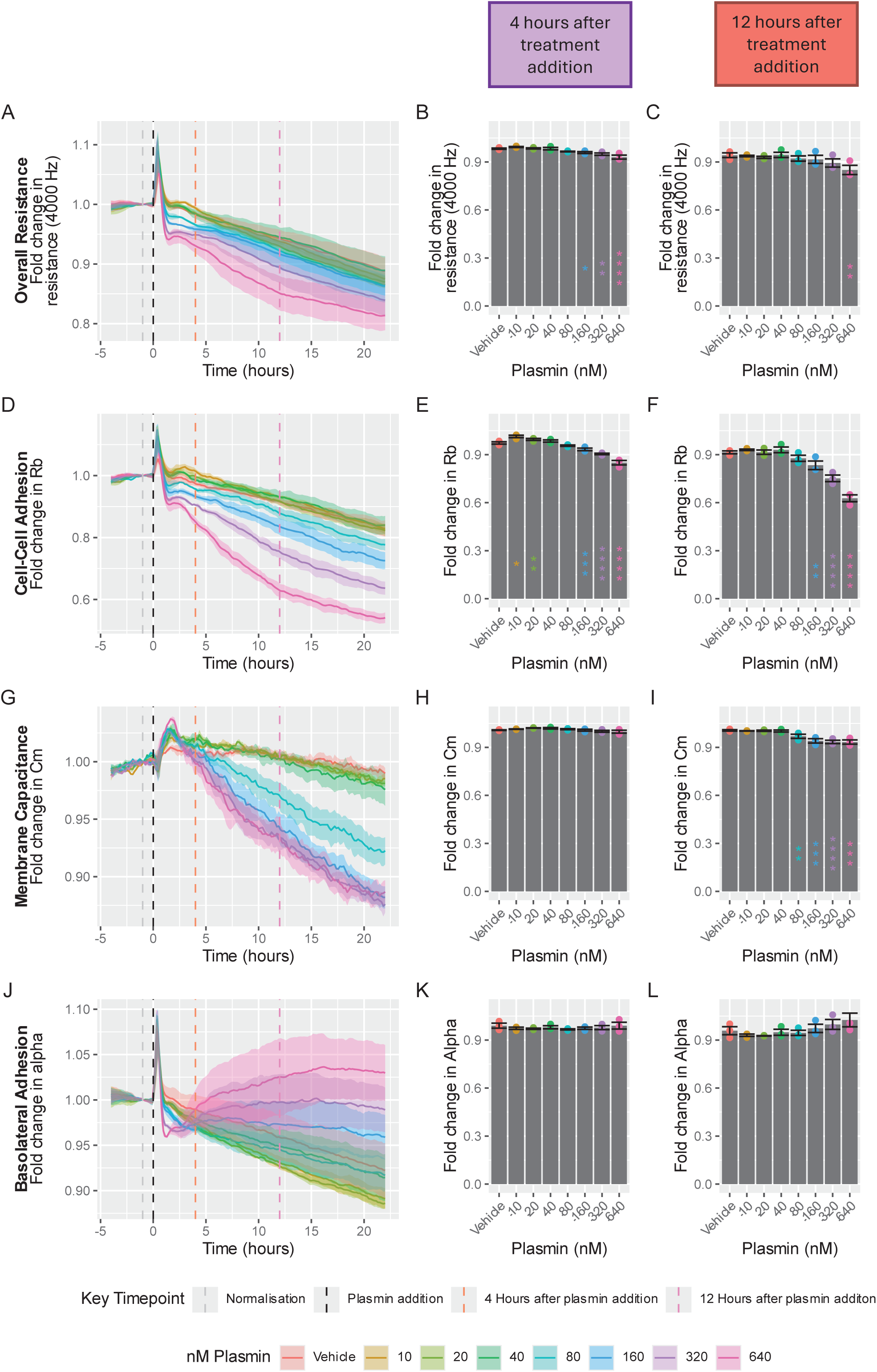
Plasmin in AIM-V-2MV serum-free media impairs the hCMEC/D3 brain endothelial barrier in a concentration dependant manner, by reducing cell to cell interactions. hCMEC/D3 cells were seeded into ECIS 96W20IDF plates and cultured in EBM-2MV serum containing media for 48 hours to allow a stable barrier to form. Cells were then washed thrice with AIM-V-2MV serum-free media and cultured in this media for 16 hours. Media was removed and replaced with respective treatment in AIM-V-2MV serum-free media. Cells were cultured for 24 hours. Barrier properties were monitored in real-time throughout the experiment using ECIS. Ribbon plots represent the mean ± SEM of data from three independent experiments, each consisting of three technical replicates (A-D). Data were normalised 2 hours before the addition of treatment using vascr. Statistical analysis of this data was carried out at 4 and 12 hours after treatment addition; each treatment response was compared with vehicle control using two-way ANOVA with Dunnett’s test (* p <0.05, ** p<0.01, *** p < 0.001,, **** p < 0.0001) (E-H).

The loss of barrier integrity was mainly attributable to plasmin causing a concentration-dependent reduction in cell-to-cell interactions (R*_b_*), with statistically significant decreases observed at higher concentrations (≥160 nM) as early as 4 hours and becoming more pronounced by 12 hours (Fig. 3D-F). In contrast, lower plasmin concentrations produced little to no change in R*_b_* compared with vehicle controls. The R*_b_* response mirrored the resistance response and was multiphasic; following treatment with ≥160 nM plasmin there was an initial rapid decline for 1 hour, which then plateaued for 2-3 hours, and subsequently declined (Fig. 3D). Plasmin also caused a small concentration-dependent decline in cell membrane capacitance (C*_m_*), with statistically significant decreases observed at higher plasmin concentrations (≥80 nM) by 12 hours after treatment (Fig. 3G-I). In contrast, lower plasmin concentrations (10-40 nM) produced little to no change in C*_m_* compared with vehicle controls. Changes in basolateral adhesion (α) were modest but multiphasic, with the highest plasmin concentrations (≥320nM) causing a rapid decline during the 1^st^ hour, which was maintained for a further ∼2-3 hours, after which the cell substrate interactions increased above the vehicle for the remainder of the experiment (Fig. 3J-L). The lower concentrations of plasmin (10-80 nM) caused a small immediate decline in basolateral adhesion during the 1^st^ hour, which then tracked slightly below the vehicle for the remainder of the experiment (Fig. 3J). Additional ECIS experiments confirmed that these responses were conserved across brain endothelial models, with plasmin also stimulating a concentration-dependent loss of hCMVEC barrier integrity driven predominantly by disruption of cell-to-cell junctions (Sup. Fig. 5).

Collectively, these data demonstrate that plasmin impaired the brain endothelial barrier in a concentration-dependent manner, principally through disruption of cell-to-cell interactions. Interestingly the response appeared multiphasic, with early rapid barrier weakening, a transient plateau, then after 4 hours high plasmin levels drove sustained barrier deterioration. To a lesser extent, plasmin also influenced membrane capacitance and basolateral adhesion of cells to the underlying substrate.

### Inhibition of plasmin activity with a2-antiplasmin prevents plasmin-induced hCMEC/D3 endothelial barrier disruption

To determine whether the barrier-disruptive effects of plasmin are dependent on its proteolytic activity, hCMEC/D3 monolayers cultured in AIM-V-2MV serum-free media were treated with plasmin in the presence or absence of the plasmin inhibitor α2-antiplasmin, and barrier properties were monitored in real-time using ECIS (Fig. 4). Consistent with previous observations, exposure to 640 nM plasmin alone caused a progressive reduction in overall barrier resistance, cell-to-cell interactions (R*_b_*) and membrane capacitance (C*_m_*), and an increase in basolateral adhesion (α). In contrast, co-treatment with equimolar α2-antiplasmin prevented these plasmin-induced changes, with each of the barrier property profiles comparable to the vehicle-treated controls (Fig. 4A, D, G & J)). Furthermore, statistical analysis confirmed that α2-antiplasmin significantly prevented plasmin-induced reductions in overall resistance, cell-to-cell interactions, and membrane capacitance at 12 hours (Fig. 4A-I). α2-antiplasmin alone did not significantly influence hCMEC/D3 endothelial barrier properties, although relatively small initial temporal elevations in resistance and R*_b_* were evident (Fig. 4A & D). Together, these data demonstrate that the deleterious effects of plasmin on the hCMEC/D3 brain endothelial barrier are dependent on its proteolytic activity because they are inhibited by α2-antiplasmin.

**Figure 4.**
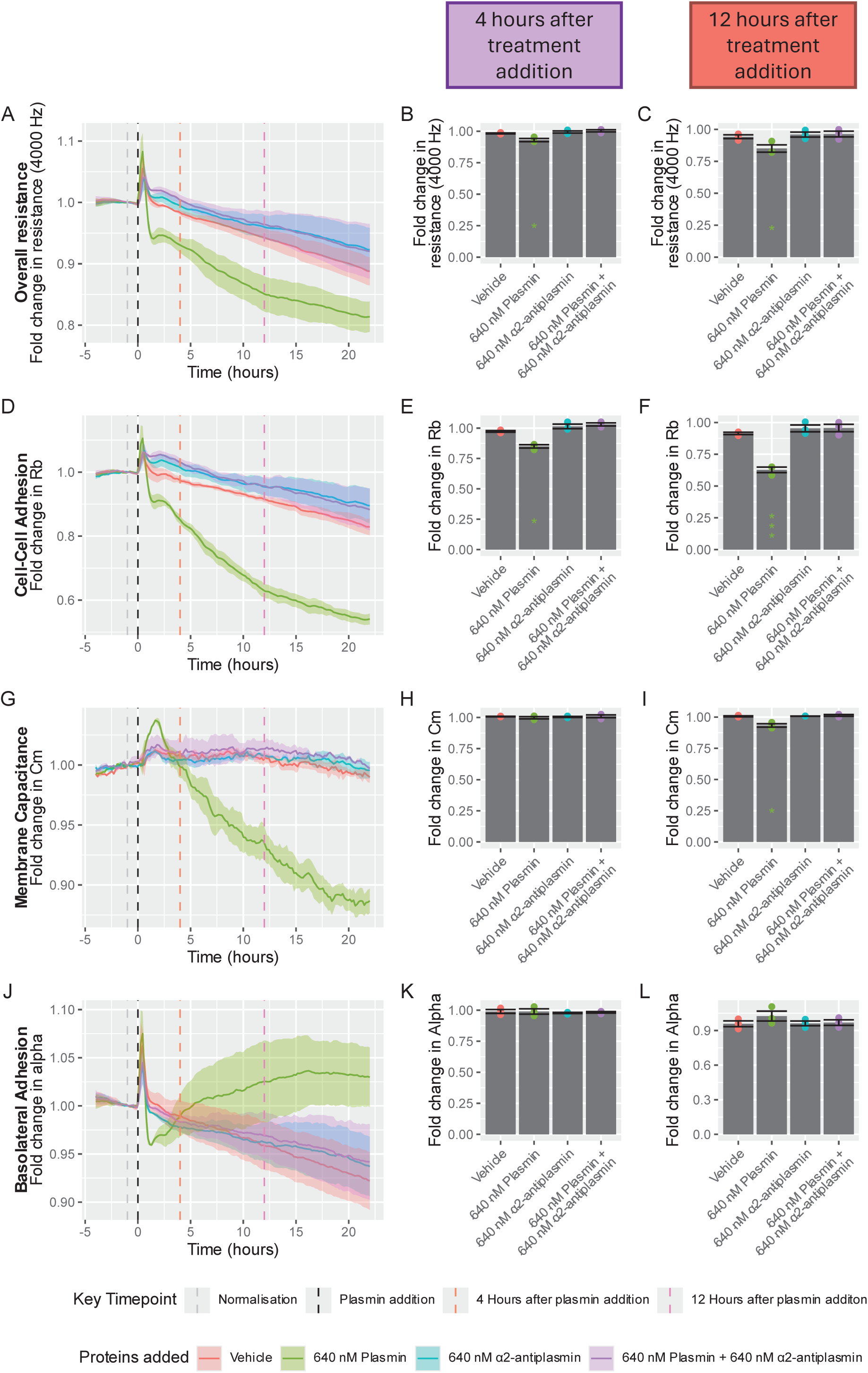
Inhibition of plasmin activity with α-2-antiplasmin prevents the plasmin induced decline of the hCMEC/D3 endothelial barrier. hCMEC/D3 cells were seeded into ECIS 96W20IDF plates and cultured in EBM-2MV serum containing media for 48 hours to allow a stable barrier to form. Cells were then washed thrice with AIM-V-2MV serum-free media and cultured in this media for 16 hours. Media was removed and replaced with respective treatment in AIM-V-2MV serum-free media. Cells were cultured for 24 hours. Barrier properties were monitored in real-time throughout the experiment using ECIS. Ribbon plots represent the mean ± SEM of data from three independent experiments, each consisting of three technical replicates (A-D). Data was normalised 1 hour before the addition of treatment using vascr. Statistical analysis of this data was carried out at 4 and 12 hours after treatment addition; each treatment response was compared with vehicle control using two-way ANOVA with Dunnett’s test (* p <0.05, *** p < 0.001).

### Plasmin reduces expression of key brain endothelial junctional proteins

To enable objective, automated quantification of junctional protein expression in large immunocytochemistry confocal image datasets we developed vjunctr, a custom image-processing workflow (Sup. Fig. 6). We then used vjunctr to perform high-throughput quantitative immunocytochemistry to assess whether plasmin-mediated barrier disruption in hCMEC/D3 monolayers cultured in AIM-V-2MV serum-free media is associated with altered junctional protein expression and localisation (Fig. 5). Cells were treated with plasmin for 4 or 12 hours and expression of principal brain endothelial junction-associated molecules were determined i.e. VE-cadherin, β-catenin, Claudin-5, ZO-1, and PECAM-1 (Fig. 5). The time points 4 and 12 hours were selected to capture key points of the multiphasic plasmin response observed with ECIS. Qualitative inspection of maximum-intensity projections reveals clearly visible changes for some of the junctional markers following plasmin treatment (Fig. 5A). Next vjunctr was used to quantify total cellular fluorescence (Fig. 5B) and the fluorescence in the junctional area (θ) (Fig. 5C). Plasmin treatment significantly reduced the total fluorescence intensity of all the junctional proteins assessed at 4 and 12 hours, with Claudin-5 exhibiting the largest decline in total expression (Fig. 5B). All junctional molecules also exhibited a significant reduction in expression in the junctional area following 4 and 12 hours of plasmin treatment, with β-catenin, Claudin-5 and PECAM-1 (CD31) displaying the largest declines (Fig. 5C).

**Figure 5.**
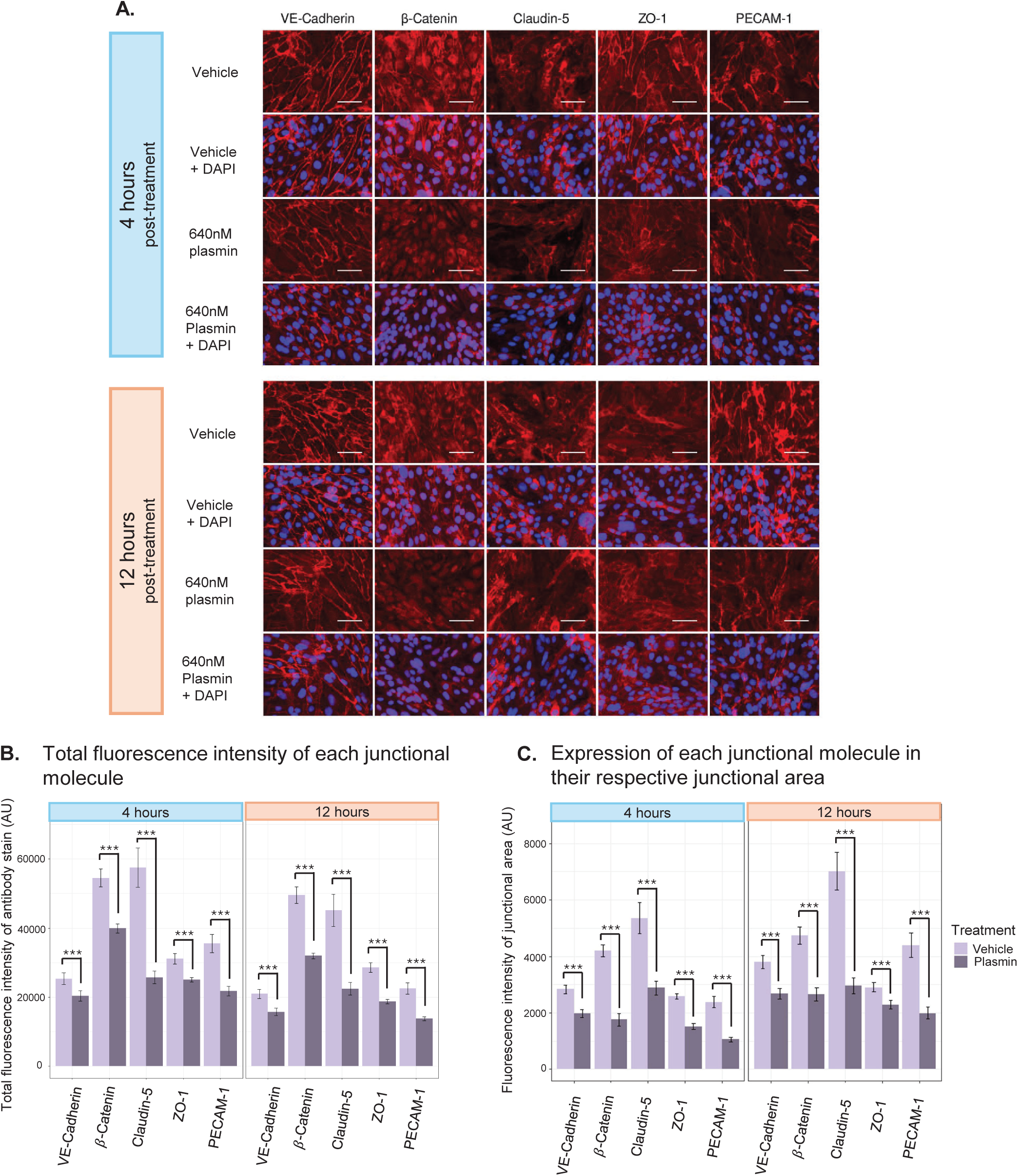
High throughput quantitative immunocytochemistry demonstrates that plasmin stimulates a decline in key brain endothelial junctional molecules. hCMEC/D3 cells were seeded into Ibidi 96 well plates in EBM-2MV serum containing media for 48 hours. Cells were then washed thrice with AIM-V-2MV serum-free media and cultured in AIM-V-2MV serum-free media for 16 hours. Vehicle or 640nM plasmin was then added and cells were cultured for 4 or 12 hours. Cells were fixed with paraformaldehyde, probed with antibodies detecting junctional molecules (VE-cadherin *(CD144),* β-Catenin, Claudin-5, ZO-1 *and PECAM-1 (CD31)*), washed, and detected with their respective fluorophore conjugated secondary antibodies and DAPI. Single planes of the wells were imaged using an Operetta CLS. The images shown are representative of 9 fields from each of three independent experiments (A). Vjunctr was then used to quantify the total fluorescence intensity and the fluorescence intensity in the junctional area for each junctional molecule (B & C). The bar charts represent the mean ± SEM of data from three independent experiments, each of which consisted of three technical replicates of nine randomly selected fields from each well. Significant differences between plasmin and vehicle control for each junctional molecule was analysed using two-way ANOVA with Tukey’s range test (*** p<0.001).

Together, these data demonstrate that plasmin stimulates a reduction in both the total expression and junctional localisation of key brain endothelial junctional proteins, providing a molecular correlate for the plasmin-mediated impairment of brain endothelial barrier integrity observed using ECIS.

## DISCUSSION

Maintaining brain microvascular endothelial cell barrier integrity is a critical determinant of outcome following ischemic stroke [26]. Loss of this barrier permits the entry of blood-derived proteins and immune cells into the brain parenchyma, driving oedema, inflammation, and secondary tissue injury [27,28]. This likely explains why BBB breakdown is strongly associated with poor neurological outcomes and increased complication rates following ischemic stroke and thrombolytic therapy [26,29,30].

While plasmin is essential for dissociating the clot during ischemic stroke, understanding how to achieve this safely is critical for improving ischemic stroke therapeutics. While both tPA and plasmin have been implicated in BBB dysfunction, distinguishing their individual roles has been challenging due to their obligatory co-occurrence *in vivo* [13]. This distinction is important therapeutically: if tPA itself compromises barrier integrity, alternative or modified plasminogen activators may be preferable. Whereas, if plasmin is the dominant effector, strategies should focus on limiting endothelial exposure to plasmin or preventing destructive downstream pathways, while preserving sufficient fibrinolytic activity [13,31,32]. This study therefore focussed on determining the direct effects of tPA and plasmin in isolation on human brain endothelial barrier integrity.

*In vivo*, BBB integrity is governed by dynamic interactions between endothelial cells, astrocytes, pericytes, neurons, and immune cells, all of which contribute to barrier maintenance and breakdown during ischemic stroke [33]. While exclusion of these supporting cell types in this study limits physiological complexity, it was necessary to establish the direct influence of plasminogen system components on brain microvascular endothelial cells. Equally critical was identifying a serum-free brain endothelial culture system; which greatly simplified data interpretation by eliminating the vast array of proteins present in serum which interact with plasminogen system proteins and alter endothelial barrier integrity [15,34–36]. Furthermore, we confirmed that following transition into serum-free conditions, the brain endothelial cells lacked detectable surface-bound plasminogen; the possibility of cells retaining plasminogen on their surface has hindered interpretation of earlier findings [15,23,34,37,38]. Collectively, our model provides unprecedented control of plasminogen system components, and ensures effects can be solely attributed to the exogenously applied plasminogen system components.

Interestingly, tPA alone did not directly disrupt brain endothelial barrier integrity. This is consistent with previous findings demonstrating no effect of tPA on barrier properties when plasmin generation is eliminated [34,39]. Although, our observation does contrast with other studies that report tPA-associated effects on endothelial barrier function; this inconsistency may reflect the generation of plasmin from residual cell surface-bound plasminogen following transfer to serum-free conditions. In these earlier studies residual surface-bound plasminogen was not quantified and it’s presence may have confounded experimental interpretation [15,34,37,38]. Importantly, tPA’s plasmin independent effects could also be mediated by other cells within the neurovascular unit, including astrocytes, pericytes and neurons, which can subsequently impair brain endothelial barrier function [15,30,31,40–42]. Together these observations suggest tPA impairs the BBB indirectly via interactions with other neurovascular unit cell types, rather than directly impacting the brain endothelial barrier.

Consistent with earlier brain and non-brain endothelial studies [43–45], our serum-free brain endothelial model exhibited barrier deterioration when cultured with both tPA and plasminogen. While plasmin generated from this reaction is likely a major contributor to barrier disruption, the concurrent presence of tPA in these assays has complicated our ability to distinguish the relative effects of these two proteases [13]. We therefore addressed this using our serum-free model and demonstrated that plasmin alone can disrupt human brain microvascular endothelial barrier integrity. This observation does differ with brain endothelial-astrocyte co-culture studies, where minimal responses to exogenous plasmin were observed. However this might reflect recognised astrocyte-mediated protective effects, or indicate the presence of residual serum-derived plasminogen system modulators in these earlier studies [15,34,37,38]. Our subsequent experiments showed prevention of barrier loss by α2-antiplasmin, confirming that plasmin directly mediated the barrier deterioration we observed. This aligns with earlier studies where α2-antiplasmin ameliorated BBB deterioration by tPA activated plasminogen, indicating plasmin contributed to this response [13,34,43]. Despite these findings, it is still not clear how plasmin mediates this affect, is it caused by plasmin’s proteolytic capacity, or a signalling pathway triggered by a cleavage product? Future studies using selective catalytic inhibitors, microplasmin that lacks the cell-surface binding domain, or multi-omics analysis may help reveal these mechanisms. It is also worth noting that although α2-antiplasmin was protective in this *in vitro* scenario, it’s role *in vivo* is complex, as elevated circulating α2-antiplasmin can worsen stroke outcomes by preventing plasmin-mediated clot breakdown & delaying reperfusion [46]; so future studies might consider specifically inhibiting plasmin localised at the endothelial surface to protect the barrier [1,13].

A strength of this study was the use of real-time ECIS, which allowed different barrier properties to be assessed for the entire experiment; this would not be possible using conventional end point permeability or lower-resolution TEER assays [17]. Modelling of the ECIS data revealed that the plasmin-induced barrier decline was predominantly caused by a reduction in the paracellular-interactions between endothelial cells. ECIS also revealed that this barrier disruption was multiphasic; there was an initial barrier decline, followed by a ∼2-hour plateaux, then a sustained gradual decline. These distinct phases indicate that this plasmin response could involve multiple mechanisms, as plasmin is promiscuous [23,39,47]. Perhaps the delayed barrier deterioration phase involves plasmin-induced activation of endothelial mediators such as matrix metalloproteinases and/or inflammatory cytokines via NF-κβ–dependent pathways [19], which previous studies suggest can be activated by plasmin [47,48]. Intriguingly, the plateau period aligns with the established ∼4-hour therapeutic window for tPA [2], and the higher levels of plasmin drive barrier deterioration beyond this point. This hints that plasmin could be a contributor to the adverse BBB effects observed beyond the tPA treatment window.

The temporal ECIS data enabled key time-points during the plasmin mediated response to be identified for subsequent immunocytochemistry analyses. Plasmin stimulated significant coordinated degradation, downregulation and/or redistribution of multiple junctional proteins, including VE-cadherin, Claudin-5, β-catenin, PECAM-1, and ZO-1. Interestingly this didn’t substantially disrupt overall barrier function, with >80% of barrier resistance (measured by ECIS) remaining at 12 hours after plasmin exposure. However, the reduction in junctional expression did correlate with the more pronounced decline in cell-to-cell interactions detected by ECIS. These observations correlate with *in vivo* stroke studies where junctional Claudin-5, VE-cadherin, and β-catenin were also downregulated [49,50]. Interestingly, β-Catenin expression at the junction declined more than total cellular expression at 4 hours, indicating redistribution to or preferable retention in other cellular regions. The images illustrate that β-catenin is predominantly located in the nucleus 4 hours after plasmin treatment, where it can act as a transcriptional regulator [51,52]. Furthermore, translocation of β-catenin away from the membrane can destabilise junctional VE-Cadherin, which in turn can disrupt junctional Claudin-5 [53,54]. Hence, perhaps β-catenin translocation represents a previously underappreciated downstream effect of plasmin on the BBB. Moreover, this may contribute to the multiphasic plasmin-response observed, and possibly the staged pathology of ischemic stroke [55]. The temporal dynamics of this plasmin response warrant further investigation, as it could reveal key-time points to protect the endothelial integrity during tPA treatment; thereby potentially extending the safe window for thrombolytic treatment.

The plasmin concentrations used in this study are physiologically relevant, aligning with recent measurements of active plasmin in infarcted murine brain and theoretical estimates during acute stroke, particularly at the endothelial surface where plasminogen is predominantly activated and inhibitors are less effective [35,56,57]. It has been argued that plasmin cannot meaningfully contribute to BBB disruption during ischemic stroke due to its efficient inhibition by α2-antiplasmin, however this can’t be entirely true as the clinical efficacy of tPA therapy necessitates plasmin-mediated fibrinolysis [13]. In fact, suppression of plasmin activity or ablation of plasminogen or tPA worsens long-term stroke outcomes by impairing clot dissolution and reperfusion [46,58–61]. Furthermore, high plasminogen levels are protective against cerebral infarction [62], underscoring the necessity of plasmin for resolving stroke. Therefore, complete inhibition of plasmin activity is not a feasible therapeutic strategy to protect the BBB during tPA treatment of ischemic stroke. Possible approaches could be to spatially restrict plasmin activation using clot targeted thrombolytics, or improving plasmin-antiplasmin balance to reduce endothelial exposure to proteolysis [63–68]. Alternatively the pathways that mediate the plasmin induced BBB disruption could be temporally interrupted downstream of proteolysis; thereby protecting the BBB whilst maintaining the plasmin activity required for clot clearance and recanalization [4].

## Conclusions

In conclusion, the data presented from this study demonstrate that plasmin can directly stimulate a reduction in brain endothelial barrier integrity, which is mediated by a decline in paracellular junctional proteins. A strength of this study was the use of a robustly validated serum-free culture model that facilitated the study of plasmin in isolation, without the influence of other plasminogen system components that would complicate attributing barrier effects specifically to plasmin.

ECIS real-time monitoring revealed that plasmin-induced barrier compromise was multiphasic. Interestingly, the 4-hr plateau phase aligned with the established therapeutic window for tPA, whilst the sustained plasmin-induced decline in barrier beyond 4 hours correlated with an increasing risk of adverse effects from thrombolysis [2,3,5,7]. Whilst this may suggest two temporally separated mechanisms are at play, these data should be interpreted within the limitations of an endothelial monoculture model, lacking various other cell types within the NVU which support BBB integrity. Collectively these data indicate that protecting the brain endothelium from plasmin could represent a potential therapeutic strategy to improve patient outcomes during the thrombolytic treatment of ischemic stroke, thereby potentially allowing extension of the thrombolytic treatment window.

## Author Contributions

The following authors contributed to study design: J. Hucklesby, C. Gao., E. Graham, and C. Angel. The following authors performed experiments: J. Hucklesby and C. Gao. The following authors contributed to the analysis and interpretation of data: J. Hucklesby, C. Gao., E. Graham, and C. Angel. The following authors contributed to writing and/or revising the intellectual content: J. Hucklesby, C. Gao, E. Graham, and C. Angel. The final draft of the manuscript was approved by J. Hucklesby, C. Gao, E. Graham, and C. Angel.

## Funding

This work was supported by the Auckland Medical Research Foundation, NZ (1217002, 2018-22), the Neurological Foundation, NZ (2330 FFE, 2023-25) and the University of Auckland, NZ (3719918, 2019-22 & PhD Scholarship awarded to C. Gao, 2025).

## Acknowledgements

We thank Jo Dodd for her generous guidance and technical support.

## Declaration of Competing Interests

J. Hucklesby, C. Gao., E. Graham, and C. Angel declare that there are no conflicts of interest.

## Data availability

For original data, please contact

## Notes

### Competing Interest Statement

The authors have declared no competing interest.

## References

1 Medcalf RL. Plasminogen and stroke: more is better. Journal of Thrombosis and Haemostasis 2016; 14: 1819–21. 10.1111/jth.13399

2 Powers WJ, Rabinstein AA, Ackerson T, Adeoye OM, Bambakidis NC, Becker K, Biller J, Brown M, Demaerschalk BM, Hoh B, Jauch EC, Kidwell CS, Leslie-Mazwi TM, Ovbiagele B, Scott PA, Sheth KN, Southerland AM, Summers DV, Tirschwell DL, on behalf of the American Heart Association Stroke Council. Guidelines for the Early Management of Patients With Acute Ischemic Stroke: 2019 Update to the 2018 Guidelines for the Early Management of Acute Ischemic Stroke: A Guideline for Healthcare Professionals From the American Heart Association/American Stroke Association. Stroke 2019; 50. 10.1161/STR.0000000000000211

3 The National Institute of Neurological Disorders and Stroke rt-PA Stroke Study Group. Tissue Plasminogen Activator for Acute Ischemic Stroke. N Engl J Med Massachusetts Medical Society; 1995; 333: 1581–8. 10.1056/NEJM199512143332401

4 Zhang B, Leung L, Su EJ, Lawrence DA. PA System in the Pathogenesis of Ischemic Stroke. ATVB 2025; 45: 600–8. 10.1161/ATVBAHA.125.322422

5 Muir KW, Ford GA, Ford I, Wardlaw JM, McConnachie A, Greenlaw N, Mair G, Sprigg N, Price CI, MacLeod MJ, Dima S, Venter M, Zhang L, O’Brien E, Sanyal R, Reid J, Sztriha LK, Haider S, Whiteley WN, Kennedy J, et al. Tenecteplase versus alteplase for acute stroke within 4·5 h of onset (ATTEST-2): a randomised, parallel group, open-label trial. The Lancet Neurology 2024; 23: 1087–96. 10.1016/S1474-4422(24)00377-6

6 Plow EF, Felez J, Miles LA. Cellular Regulation of Fibrinolysis. Thromb Haemost 1991; 66: 032–6. 10.1055/s-0038-1646369

7 Hacke W, Kaste M, Bluhmki E, Brozman M, Dávalos A, Guidetti D, Larrue V, Lees KR, Medeghri Z, Machnig T, Schneider D, Von Kummer R, Wahlgren N, Toni D. Thrombolysis with Alteplase 3 to 4.5 Hours after Acute Ischemic Stroke. N Engl J Med 2008; 359: 1317–29. 10.1056/NEJMoa0804656

8 Saini V, Guada L, Yavagal DR. Global Epidemiology of Stroke and Access to Acute Ischemic Stroke Interventions. Neurology 2021; 97. 10.1212/WNL.0000000000012781

9 Liang Y, Yu Y, Liu J, Li X, Chen X, Zhou H, Guo Z-N. Blood-brain barrier disruption and hemorrhagic transformation in acute stroke before endovascular reperfusion therapy. Front Neurol 2024; 15: 1349369. 10.3389/fneur.2024.1349369

10 Rust R, Yin H, Achón Buil B, Sagare AP, Kisler K. The blood–brain barrier: a help and a hindrance. Brain 2025; 148: 2262–82. 10.1093/brain/awaf068

11 Kadry H, Noorani B, Cucullo L. A blood–brain barrier overview on structure, function, impairment, and biomarkers of integrity. Fluids Barriers CNS 2020; 17: 69. 10.1186/s12987-020-00230-3

12 Privratsky JR, Newman PJ. PECAM-1: regulator of endothelial junctional integrity. Cell Tissue Res 2014; 355: 607–19. 10.1007/s00441-013-1779-3

13 Niego B, Medcalf RL. Plasmin-Dependent Modulation of the Blood–Brain Barrier: A Major Consideration during tPA-Induced Thrombolysis? J Cereb Blood Flow Metab 2014; 34: 1283–96. 10.1038/jcbfm.2014.99

14 Ishiguro M, Mishiro K, Fujiwara Y, Chen H, Izuta H, Tsuruma K, Shimazawa M, Yoshimura S, Satoh M, Iwama T, Hara H. Phosphodiesterase-III Inhibitor Prevents Hemorrhagic Transformation Induced by Focal Cerebral Ischemia in Mice Treated with tPA. Kleinschnitz C, editor. PLoS ONE 2010; 5: e15178. 10.1371/journal.pone.0015178

15 Niego B, Lee N, Larsson P, De Silva TM, Au AE-L, McCutcheon F, Medcalf RL. Selective inhibition of brain endothelial Rho-kinase-2 provides optimal protection of an in vitro blood-brain barrier from tissue-type plasminogen activator and plasmin. Wang X, editor. PLoS ONE 2017; 12: e0177332. 10.1371/journal.pone.0177332

16 Weksler BB, Subileau EA, Perrière N, Charneau P, Holloway K, Leveque M, Tricoire-Leignel H, Nicotra A, Bourdoulous S, Turowski P, Male DK, Roux F, Greenwood J, Romero IA, Couraud PO. Blood-brain barrier-specific properties of a human adult brain endothelial cell line. The FASEB Journal 2005; 19: 1872–4. 10.1096/fj.04-3458fje

17 Hucklesby JJW, Anchan A, O’Carroll SJ, Unsworth CP, Graham ES, Angel CE. Comparison of Leading Biosensor Technologies to Detect Changes in Human Endothelial Barrier Properties in Response to Pro-Inflammatory TNFa and IL1β in Real-Time. Biosensors 2021; 11: 159. 10.3390/bios11050159

18 Robilliard LD, Kho DT, Johnson RH, Anchan A, O’Carroll SJ, Graham ES. The Importance of Multifrequency Impedance Sensing of Endothelial Barrier Formation Using ECIS Technology for the Generation of a Strong and Durable Paracellular Barrier. Biosensors 2018; 8: 64. 10.3390/bios8030064

19 O’Carroll SJ, Kho DT, Wiltshire R, Nelson V, Rotimi O, Johnson R, Angel CE, Graham ES. Pro-inflammatory TNFa and IL-1β differentially regulate the inflammatory phenotype of brain microvascular endothelial cells. J Neuroinflammation 2015; 12: 131. 10.1186/s12974-015-0346-0

20 Wickham H. ggplot2. Cham: Springer International Publishing; 2016. 10.1007/978-3-319-24277-4

21 Wegener J, Keese CR, Giaever I. Electric Cell–Substrate Impedance Sensing (ECIS) as a Noninvasive Means to Monitor the Kinetics of Cell Spreading to Artificial Surfaces. Experimental Cell Research 2000; 259: 158–66. 10.1006/excr.2000.4919

22 Hucklesby JJW, Unsworth CP, Graham ES, Angel CE. vascr: An R-based toolkit for rapid and robust analysis of cellular impedance sensing data. Cell Biology; 2025. 10.64898/2025.12.17.695026

23 Hucklesby JJW, Angel CE, Graham ES, Dunbar PR, Birch NP, Loef EJ. Plasmin reduces human T cell arrest on endothelial-like cells by cleaving bound CCL21 from the cell surface. Experimental Cell Research 2025; 446: 114480. 10.1016/j.yexcr.2025.114480

24 Link AJ, LaBaer J. Proteomics: A Cold Spring Harbor Laboratory Course Manual. Cold Spring Harbor Laboratory Press; 2009.

25 Stief TW, Richter A, Bünder R, Maisch B, Renz H. Monitoring of Plasmin and Plasminogen Activator Activity in Blood of Patients under Fibrinolytic Treatment by Reteplase. Clin Appl Thromb Hemost 2006; 12: 213–8. 10.1177/107602960601200210

26 Nadareishvili Z, Simpkins AN, Hitomi E, Reyes D, Leigh R. Post-Stroke Blood-Brain Barrier Disruption and Poor Functional Outcome in Patients Receiving Thrombolytic Therapy. Cerebrovasc Dis 2019; 47: 135–42. 10.1159/000499666

27 Chu HX, Kim HA, Lee S, Moore JP, Chan CT, Vinh A, Gelderblom M, Arumugam TV, Broughton BR, Drummond GR, Sobey CG. Immune Cell Infiltration in Malignant Middle Cerebral Artery Infarction: Comparison with Transient Cerebral Ischemia. J Cereb Blood Flow Metab 2014; 34: 450–9. 10.1038/jcbfm.2013.217

28 Kidwell CS, Latour L, Saver JL, Alger JR, Starkman S, Duckwiler G, Jahan R, Vinuela F, Kang D-W, Warach S. Thrombolytic Toxicity: Blood Brain Barrier Disruption in Human Ischemic Stroke. Cerebrovasc Dis 2008; 25: 338–43. 10.1159/000118379

29 Luissint A-C, Artus C, Glacial F, Ganeshamoorthy K, Couraud P-O. Tight junctions at the blood brain barrier: physiological architecture and disease-associated dysregulation. Fluids Barriers CNS 2012; 9: 23. 10.1186/2045-8118-9-23

30 Sifat AE, Vaidya B, Abbruscato TJ. Blood-Brain Barrier Protection as a Therapeutic Strategy for Acute Ischemic Stroke. AAPS J 2017; 19: 957–72. 10.1208/s12248-017-0091-7

31 Kaur J, Zhao Z, Klein GM, Lo EH, Buchan AM. The Neurotoxicity of Tissue Plasminogen Activator? J Cereb Blood Flow Metab 2004; 24: 945–63. 10.1097/01.WCB.0000137868.50767.E8

32 Marder VJ, Stewart D. Towards safer thrombolytic therapy. Seminars in Hematology 2002; 39: 206–16. 10.1053/shem.2002.34088

33 Qin C, Yang S, Chu Y-H, Zhang H, Pang X-W, Chen L, Zhou L-Q, Chen M, Tian D-S, Wang W. Signaling pathways involved in ischemic stroke: molecular mechanisms and therapeutic interventions. Sig Transduct Target Ther 2022; 7: 215. 10.1038/s41392-022-01064-1

34 Niego B, Freeman R, Puschmann TB, Turnley AM, Medcalf RL. t-PA–specific modulation of a human blood-brain barrier model involves plasmin-mediated activation of the Rho kinase pathway in astrocytes. Blood 2012; 119: 4752–61. 10.1182/blood-2011-07-369512

35 Cederholm-Williams S. Concentration of plasminogen and antiplasmin in plasma and serum. J Clin Pathol 1981; 34: 979–81. 10.1136/jcp.34.9.979

36 Cesarman-Maus G, Hajjar KA. Molecular mechanisms of fibrinolysis. Br J Haematol 2005; 129: 307–21. 10.1111/j.1365-2141.2005.05444.x

37 Freeman R, Niego B, R. Croucher D, Pedersen LO, Medcalf RL. t-PA, but not desmoteplase, induces plasmin-dependent opening of a blood-brain barrier model under normoxic and ischaemic conditions. Brain Research 2014; 1565: 63–73. 10.1016/j.brainres.2014.03.027

38 Keragala CB, Woodruff TM, Liu Z, Niego B, Ho H, McQuilten Z, Medcalf RL. Tissue-Type Plasminogen Activator and Tenecteplase-Mediated Increase in Blood Brain Barrier Permeability Involves Cell Intrinsic Complement. Front Neurol 2020; 11: 577272. 10.3389/fneur.2020.577272

39 Niego B, Medcalf RL. Plasmin-Dependent Modulation of the Blood–Brain Barrier: A Major Consideration during tPA-Induced Thrombolysis? J Cereb Blood Flow Metab 2014; 34: 1283–96. 10.1038/jcbfm.2014.99

40 Fan M, Xu H, Wang L, Luo H, Zhu X, Cai P, Wei L, Lu L, Cao Y, Ye R, Fan W, Zhao B-Q. Tissue Plasminogen Activator Neurotoxicity is Neutralized by Recombinant ADAMTS 13. Sci Rep 2016; 6: 25971. 10.1038/srep25971

41 Polavarapu R, Gongora MC, Yi H, Ranganthan S, Lawrence DA, Strickland D, Yepes M. Tissue-type plasminogen activator–mediated shedding of astrocytic low-density lipoprotein receptor–related protein increases the permeability of the neurovascular unit. Blood 2007; 109: 3270–8. 10.1182/blood-2006-08-043125

42 Yang E, Cai Y, Yao X, Liu J, Wang Q, Jin W, Wu Q, Fan W, Qiu L, Kang C, Wu J. Tissue plasminogen activator disrupts the blood-brain barrier through increasing the inflammatory response mediated by pericytes after cerebral ischemia. Aging (Albany NY) 2019; 11: 10167–82. 10.18632/aging.102431

43 Nagy Z, Kolev K, Csonka É, Pék M, Machovich R. Contraction of Human Brain Endothelial Cells Induced by Thrombogenic and Fibrinolytic Factors: An In Vitro Cell Culture Model. Stroke 1995; 26: 265–70. 10.1161/01.STR.26.2.265

44 Conforti G, Dominguez-Jimenez C, Ronne E, Hoyer-Hansen G, Dejana E. Cell-surface plasminogen activation causes a retraction of in vitro cultured human umbilical vein endothelial cell monolayer. Blood 1994; 83: 994–1005. 10.1182/blood.V83.4.994.994

45 Davis GE, Allen KAP, Salazar R, Maxwell SA. Matrix metalloproteinase-1 and −9 activation by plasmin regulates a novel endothelial cell-mediated mechanism of collagen gel contraction and capillary tube regression in three-dimensional collagen matrices. Journal of Cell Science 2001; 114: 917–30. 10.1242/jcs.114.5.917

46 Singh S, Saleem S, Reed GL. Alpha2-Antiplasmin: The Devil You Don’t Know in Cerebrovascular and Cardiovascular Disease. Front Cardiovasc Med 2020; 7: 608899. 10.3389/fcvm.2020.608899

47 Syrovets T, Lunov O, Simmet T. Plasmin as a proinflammatory cell activator. Journal of Leukocyte Biology 2012; 92: 509–19. 10.1189/jlb.0212056

48 Draxler D, Sashindranath M, Medcalf R. Plasmin: A Modulator of Immune Function. Semin Thromb Hemost 2016; 43: 143–53. 10.1055/s-0036-1586227

49 Teng F, Beray-Berthat V, Coqueran B, Lesbats C, Kuntz M, Palmier B, Garraud M, Bedfert C, Slane N, Bérézowski V, Szeremeta F, Hachani J, Scherman D, Plotkine M, Doan B-T, Marchand-Leroux C, Margaill I. Prevention of rt-PA induced blood–brain barrier component degradation by the poly(ADP-ribose)polymerase inhibitor PJ34 after ischemic stroke in mice. Experimental Neurology 2013; 248: 416–28. 10.1016/j.expneurol.2013.07.007

50 Song S, Huang H, Guan X, Fiesler V, Bhuiyan MIH, Liu R, Jalali S, Hasan MN, Tai AK, Chattopadhyay A, Chaparala S, Sun M, Stolz DB, He P, Agalliu D, Sun D, Begum G. Activation of endothelial Wnt/β-catenin signaling by protective astrocytes repairs BBB damage in ischemic stroke. Progress in Neurobiology 2021; 199: 101963. 10.1016/j.pneurobio.2020.101963

51 Tran KA, Zhang X, Predescu D, Huang X, Machado RF, Göthert JR, Malik AB, Valyi-Nagy T, Zhao Y-Y. Endothelial β-Catenin Signaling Is Required for Maintaining Adult Blood–Brain Barrier Integrity and Central Nervous System Homeostasis. Circulation 2016; 133: 177–86. 10.1161/CIRCULATIONAHA.115.015982

52 Valenta T, Hausmann G, Basler K. The many faces and functions of β-catenin: β-Catenin: a life by, beyond, and against the Wnt canon. The EMBO Journal 2012; 31: 2714–36. 10.1038/emboj.2012.150

53 Liebner S, Corada M, Bangsow T, Babbage J, Taddei A, Czupalla CJ, Reis M, Felici A, Wolburg H, Fruttiger M, Taketo MM, Von Melchner H, Plate KH, Gerhardt H, Dejana E. Wnt/β-catenin signaling controls development of the blood–brain barrier. The Journal of Cell Biology 2008; 183: 409–17. 10.1083/jcb.200806024

54 Taddei A, Giampietro C, Conti A, Orsenigo F, Breviario F, Pirazzoli V, Potente M, Daly C, Dimmeler S, Dejana E. Endothelial adherens junctions control tight junctions by VE-cadherin-mediated upregulation of claudin-5. Nat Cell Biol 2008; 10: 923–34. 10.1038/ncb1752

55 Sandoval KE, Witt KA. Blood-brain barrier tight junction permeability and ischemic stroke. Neurobiology of Disease 2008; 32: 200–19. 10.1016/j.nbd.2008.08.005

56 Félez J. Plasminogen binding to cell surfaces. Fibrinolysis and Proteolysis 1998; 12: 183–9. 10.1016/S0268-9499(98)80012-X

57 Mindel E, Weiss R, Bushi D, Gera O, Orion D, Chapman J, Shavit-Stein E. Increased brain plasmin levels following experimental ischemic stroke in male mice. J of Neuroscience Research 2021; 99: 966–76. 10.1002/jnr.24764

58 Grocott HP, Sheng H, Miura Y, Sarraf-Yazdi S, Mackensen GB, Pearlstein RD, Warner DS. The Effects of Aprotinin on Outcome from Cerebral Ischemia in the Rat. Anesthesia & Analgesia 1999; 88: 1–7. 10.1213/00000539-199901000-00001

59 Atochin DN, Murciano JC, Gürsoy-Özdemir Y, Krasik T, Noda F, Ayata C, Dunn AK, Moskowitz MA, Huang PL, Muzykantov VR. Mouse Model of Microembolic Stroke and Reperfusion. Stroke 2004; 35: 2177–82. 10.1161/01.STR.0000137412.35700.0e

60 Nagai N, De Mol M, Lijnen HR, Carmeliet P, Collen D. Role of Plasminogen System Components in Focal Cerebral Ischemic Infarction: A Gene Targeting and Gene Transfer Study in Mice. Circulation 1999; 99: 2440–4. 10.1161/01.CIR.99.18.2440

61 Yepes M, Sandkvist M, Moore EG, Bugge TH, Strickland DK, Lawrence DA. Tissue-type plasminogen activator induces opening of the blood-brain barrier via the LDL receptor–related protein. J Clin Invest 2003; 112: 1533–40. 10.1172/JCI200319212

62 Singh S, Houng AK, Wang D, Reed GL. Physiologic variations in blood plasminogen levels affect outcomes after acute cerebral thromboembolism in mice: a pathophysiologic role for microvascular thrombosis. Journal of Thrombosis and Haemostasis 2016; 14: 1822–32. 10.1111/jth.13390

63 Spanò R, Portioli C, Geroski T, Felici A, Palange AL, Gawne PJ, Mamberti S, Avancini G, Palomba R, Bonnard T, Moore TL, Del Sette M, Filipovic N, Vivien D, Decuzzi P. Enhancing Thrombolysis Safety in Post-Acute Ischemic Stroke with Tissue Plasminogen Activator-Associated Microparticles. ACS Nano 2025; 19: 22882–99. 10.1021/acsnano.5c01499

64 Cheng J-W, Zhang X-J, Cheng L-S, Li G-Y, Zhang L-J, Ji K-X, Zhao Q, Bai Y. Low-Dose Tissue Plasminogen Activator in Acute Ischemic Stroke: A Systematic Review and Meta-Analysis. Journal of Stroke and Cerebrovascular Diseases 2018; 27: 381–90. 10.1016/j.jstrokecerebrovasdis.2017.09.014

65 Palazzolo JS, Ale A, Ho H, Jagdale S, Broughton BRS, Medcalf RL, Wright DK, Alt K, Hagemeyer CE, Niego B. Platelet-targeted thrombolysis for treatment of acute ischemic stroke. Blood Advances 2023; 7: 561–74. 10.1182/bloodadvances.2021006691

66 Disharoon D, Trewyn BG, Herson PS, Marr DWM, Neeves KB. Breaking the fibrinolytic speed limit with microwheel co-delivery of tissue plasminogen activator and plasminogen. Journal of Thrombosis and Haemostasis 2022; 20: 486–97. 10.1111/jth.15617

67 Aamir R, Fyffe C, Korin N, Lawrence DA, Su EJ, Kanapathipillai M. Heparin and Arginine Based Plasmin Nanoformulation for Ischemic Stroke Therapy. IJMS 2021; 22: 11477. 10.3390/ijms222111477

68 Yogendrakumar V, Vandelanotte S, Mistry EA, Hill MD, Coutts SB, Nogueira RG, Nguyen TN, Medcalf RL, Broderick JP, De Meyer SF, Campbell BCV. Emerging Adjuvant Thrombolytic Therapies for Acute Ischemic Stroke Reperfusion. Stroke 2024; 55: 2536–46. 10.1161/STROKEAHA.124.045755

