## Supplementary Figure 1 for "Plasmin, the product of tissue plasminogen activator (tPA) treatment for ischemic stroke, impairs human brain endothelial barrier integrity"

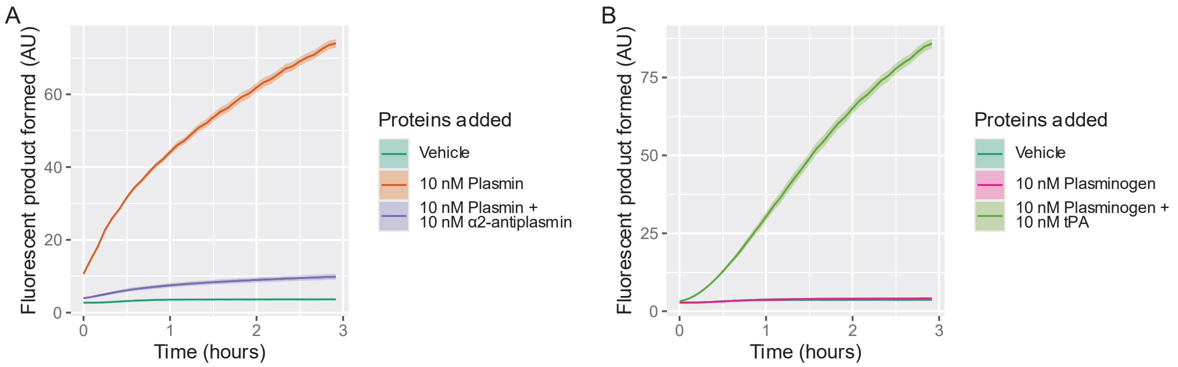


**Supplementary Figure 1. A D-VLC-AMC plasmin fluorescent substrate assay confirmed that the commercial plasmin, α2-antiplasmin, plasminogen, and tPA used for this study exhibited the expected functional activities.**

The D-VLC-AMC plasmin-specific fluorogenic substrate was rapidly cleaved by 10 nM plasmin into fluorescent product. In contrast, vehicle-treated wells exhibited minimal fluorescence, indicating low background substrate cleavage. These data confirm that the plasmin is enzymatically active (A). Co-incubation of plasmin with 10 nM α2-antiplasmin substantially attenuated plasmin-mediated substrate cleavage, reducing fluorescence levels to near-background values (A); this confirms effective inhibition of plasmin's enzymatic activity by α2-antiplasmin under these conditions (A). Plasminogen alone didn't produce a detectable increase in fluorescence above vehicle control, consistent with its inactive zymogen state (B). However, addition of 10 nM tPA to plasminogen resulted in D-VLC-AMC cleavage, reaching fluorescence levels comparable to active plasmin (B). This demonstrates efficient conversion of plasminogen into active plasmin by tPA. Collectively, these results verify that plasmin, α2-antiplasmin, plasminogen, and tPA are functionally active.

Briefly, DMEM containing 10 mM D-VLC-AMC was added to each well of a clear bottom black-sided 96 well plate. Plasmin, α2-antiplasmin, plasminogen & tPA, or their respective vehicles were then introduced. The plate was sealed and incubated at 37°C in a SpectraMax iD3 plate reader; fluorescence readings were taken every 5 minutes for 16 hours. The data presented are the mean ± SEM of three technical replicates. AU represents arbitrary units.
