## Supplementary Figure 2 for "Plasmin, the product of tissue plasminogen activator (tPA) treatment for ischemic stroke, impairs human brain endothelial barrier integrity"

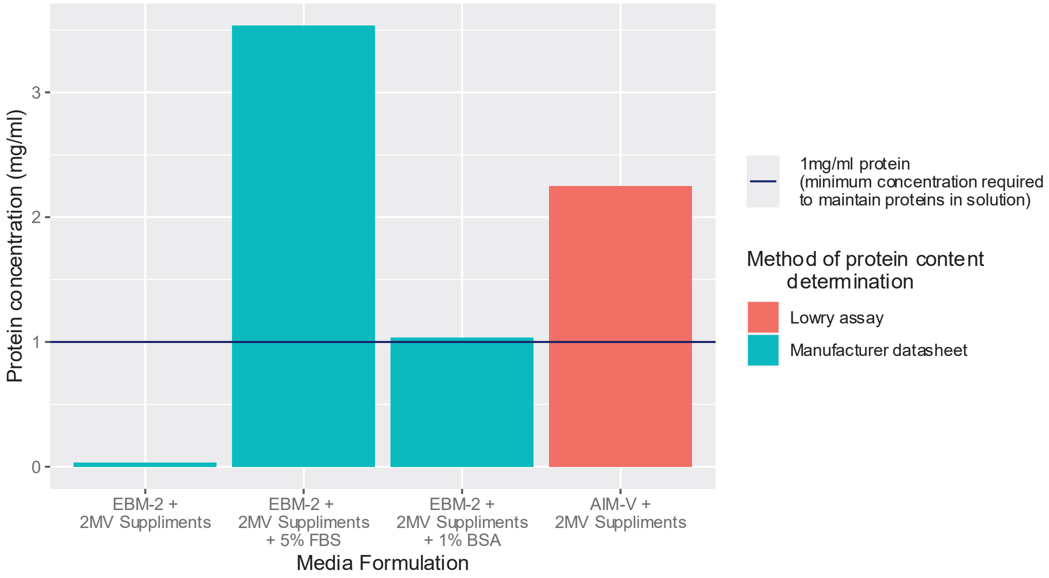


**Supplementary Figure 2. EBM-2 media + 2MV supplements contains insufficient protein to support low level protein in solution. Whilst AIM-V + 2MV supplements contains sufficient protein to support low level exogenously added protein in solution*.***

The horizontal blue line indicates 1mg/ml protein, the minimum protein concentration required to support low protein levels in solution [S1]. Apart from AIM-V media, protein concentration data was gathered from manufacturer's datasheets. The AIM-V manufacturer does not disclose this media's total protein concentration; therefore, the value presented was measured using the Lowry protein assay. Briefly, protein assays were conducted in 96 well plates, according to the manufacturer's instructions (Bio-Rad, 5000112). Absorbance readings were captured using a SpectraMax ID3 plate reader (Molecular Devices) and converted to absolute concentrations using a standard curve generated in Excel (Microsoft, V 2301).

Supplementary references

1. Link AJ, LaBaer J. Proteomics: A Cold Spring Harbor Laboratory Course Manual. Cold Spring Harbor Laboratory Press; 2009.
