## Supplementary Figure 3 for "Plasmin, the product of tissue plasminogen activator (tPA) treatment for ischemic stroke, impairs human brain endothelial barrier integrity"

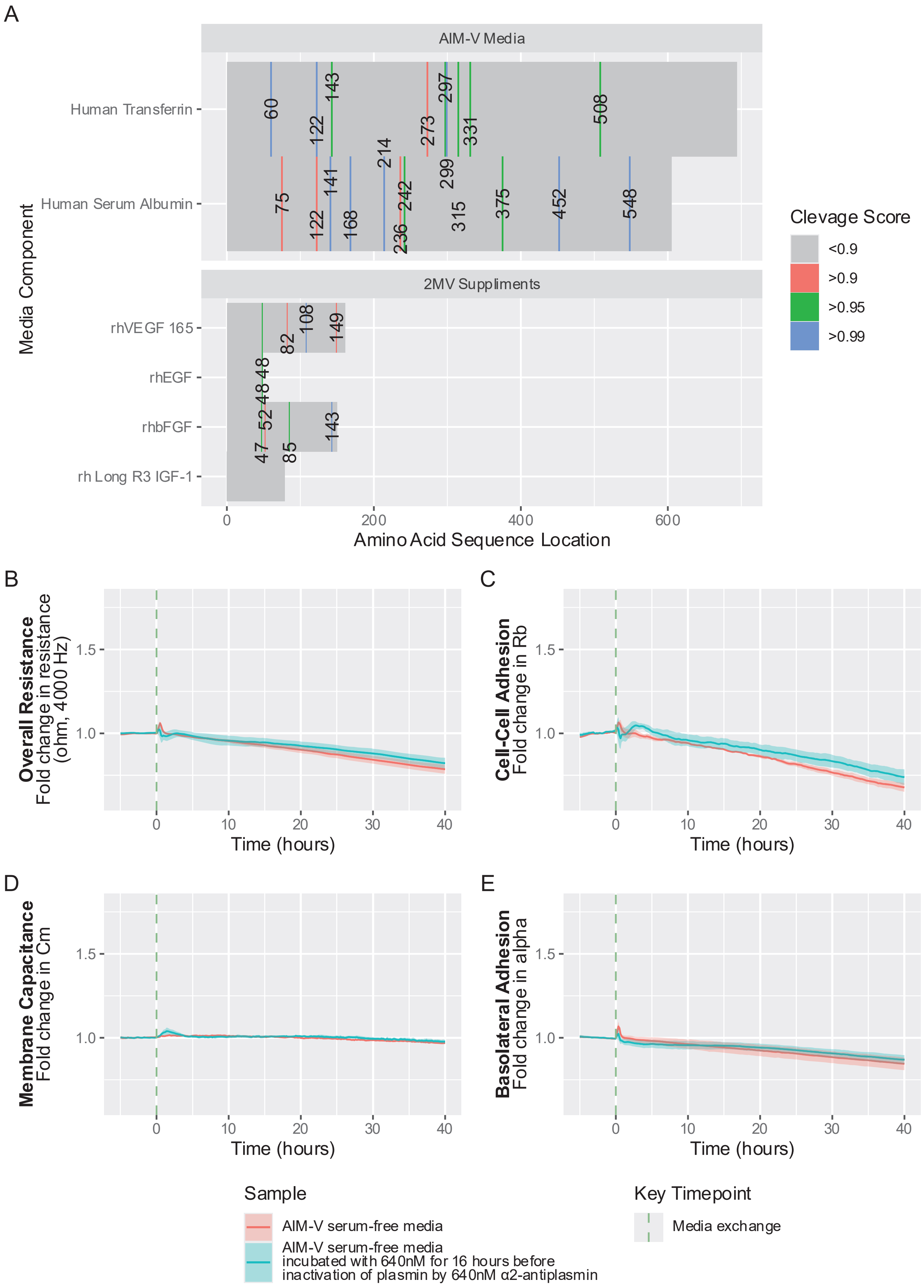

**Supplementary Figure 3. *In silico* analysis suggested that components of the AIM-V-2MV serum-free media may be susceptible to cleavage by plasmin. However, ECIS demonstrated that plasmin does not impair the ability of AIM-V-2MV serum-free media to support a stable hCMEC/D3 endothelial barrier.**

Initial *in silico* analysis suggested that components of the AIM-V-2MV serum-free media might be susceptible to cleavage by plasmin (A). Natural protein amino acid sequences were derived from the UniProt database (Release 23_01), and the recombinant 2MV supplement amino acid sequences were retrieved from Peprotech. Potential locations for plasmin cleavage were then identified using ProCleave [S2], and the resulting cleavage scores were visualised using ggplot2. A higher cleavage score indicates a higher likelihood of cleavage occurring at that site in the protein (A).

Subsequent ECIS experiments demonstrated that AIM-V-2MV serum-free media pre-cultured with plasmin retained its capacity to support a functional hCMEC/D3 brain endothelial barrier (B). Briefly, hCMEC/D3 cells were seeded into ECIS 96W20IDF plates and cultured in EBM-2MV serum containing media for 48 hours until a stable barrier formed. Cells were then washed thrice with AIM-V-2MV serum-free media. Cells were then cultured for 40 hours in either AIM-V-2MV serum-free media, or AIM-V-2MV serum-free media that had been treated with 640nM plasmin for 16 hrs and thereafter 640nM $\alpha_2$-antiplasmin to inactivate the plasmin. hCMEC/D3 barrier properties were monitored in real-time throughout the experiment using ECIS. The data presented are the mean ± SEM from three independent experiments, each consisting of three technical replicates.

Supplementary references

2. Li F, Leier A, Liu Q, Wang Y, Xiang D, Akutsu T, Webb GI, Smith AI, Marquez-Lago T, Li J, Song J. Procleave: Predicting Protease-Specific Substrate Cleavage Sites by Combining Sequence and Structural Information. *Genomics, Proteomics & Bioinformatics* 2020; **18**: 52–64.
