## Supplementary Figure 4 for "Plasmin, the product of tissue plasminogen activator (tPA) treatment for ischemic stroke, impairs human brain endothelial barrier integrity"

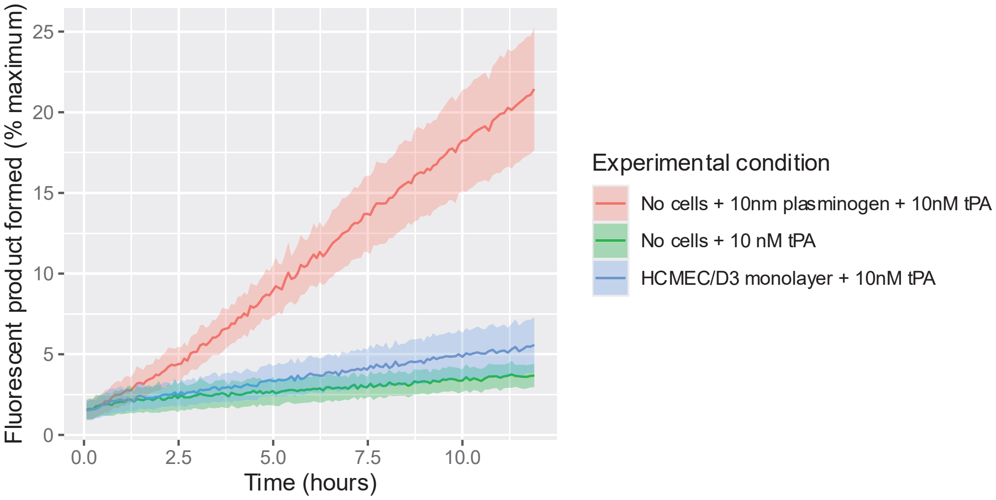


**Supplementary Figure 4. The D-VLC-AMC plasmin fluorescent substrate assay confirms that minimal plasminogen remains bound to the surface of hCMEC/D3 monolayers following transition into in AIM-V-MV2 serum-free media.**

It was important to confirm the absence of cell surface-bound plasminogen following transition from serum-containing media into the AIM-V-2MV serum-free media. This ensured that observed effects reflect plasmin treatment alone, rather than serum-derived plasminogen retained on the cell surface and later dissociating and becoming converted to plasmin, a confounding factor suspected in earlier studies [S3-5].

Importantly, data in Sup Fig 3. demonstrates that the hCMEC/D3 brain endothelial barrier is stable in AIM-V-2MV serum-free media, hence we were able to incorporate rigorous wash-steps and a 16-hour culture period to optimise dissociation of serum derived plasminogen from the cell surface. D-VLC-AMC plasmin-specific fluorogenic substrate assays, confirmed that hCMEC/D3 monolayers transitioned into AIM-V-2MV serum-free media and treated with tPA, exhibited minimal plasmin activity (A). Therefore, hCMEC/D3 brain endothelial cells transitioned from EBM-2MV serum-containing media to AIM-V-2MV serum-free media retained minimal cell surface bound plasminogen.

Briefly, cells were seeded into clear bottom black-sided 96 well plates and cultured in EBM 2MV serum-containing media for 48 hours. Cells were then washed thrice with AIM-V-2MV serum-free media and cultured in this media for 16 hours. Media was removed and replaced with respective treatment in AIM-V-2MV serum-free media with 10 mM D-VLC-AMC plasmin fluorescent substrate. The plate was sealed and incubated at 37°C in a SpectraMax iD3 plate reader; fluorescence readings were taken every 5 minutes for 12 hours. The data presented are the mean ± SEM of three technical replicates. 100% represents cleavage by 100nM plasmin at 16hr.
