## Supplementary Figure 5 for "Plasmin, the product of tissue plasminogen activator (tPA) treatment for ischemic stroke, impairs human brain endothelial barrier integrity"

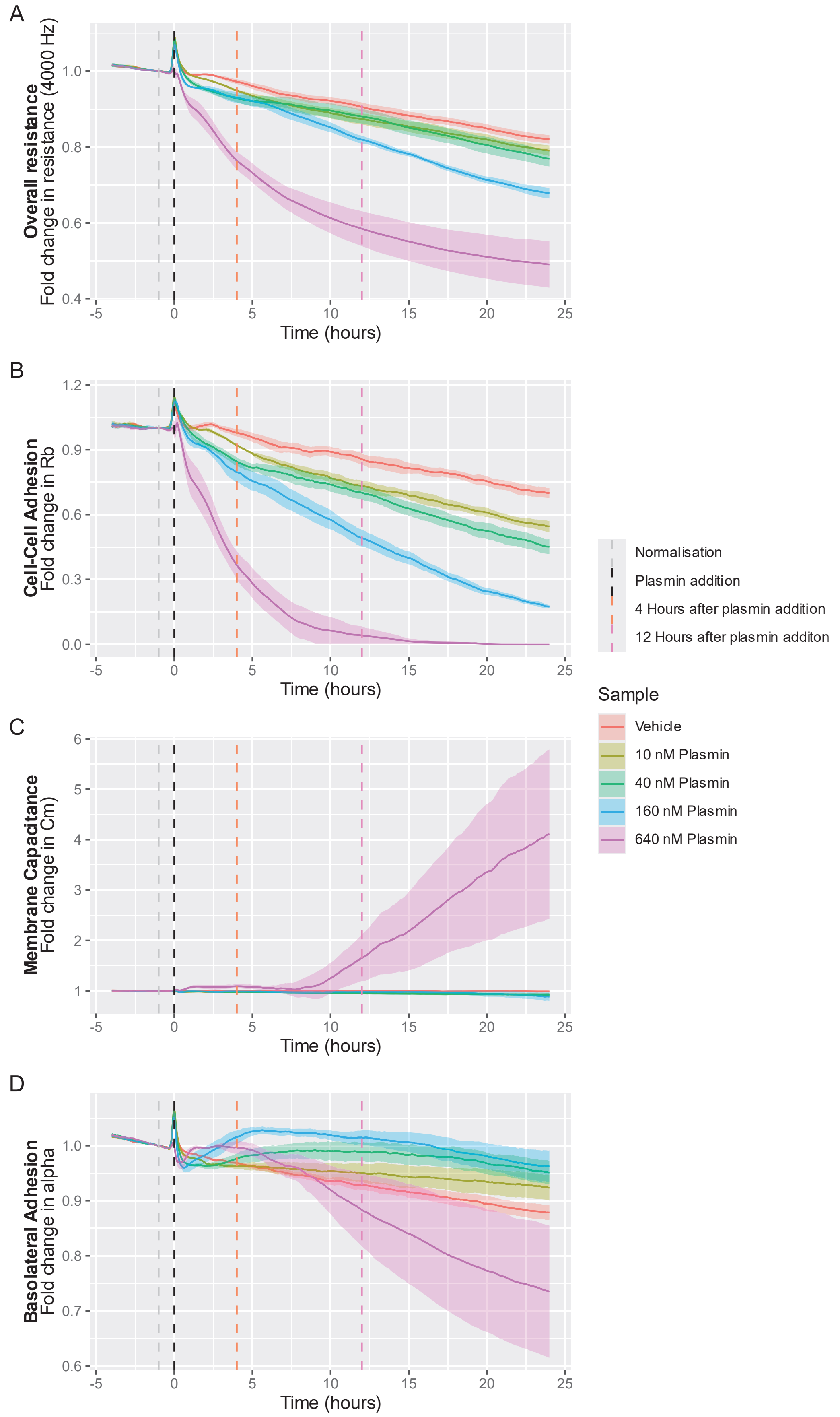

**Supplementary Figure 5. Plasmin in AIM-V-hCMVEC supplement serum-free media impairs the hCMVEC brain endothelial barrier in a concentration dependent manner, by reducing cell-to-cell interactions.**

Plasmin caused a progressive, concentration-dependent decline in hCMVEC barrier resistance over 24 hours, with more pronounced effects at higher concentrations (≥160 nM) and minimal effects at lower concentrations (A). ECIS modelling indicates that barrier disruption was driven primarily by impaired cell-to-cell interactions, with complete loss of $R_b$ with 640nM plasmin, and concentration-dependent reductions with 10-160 nM plasmin by 12 hours (B). Incubation with 640nM plasmin also caused membrane capacitance ($C_m$) to increase substantially and basolateral adhesion (alpha) to decline (C & D), consistent with the junctional destabilisation and loss of barrier integrity observed. Plasmin concentration (10-160 nM) dependant increases in basolateral adhesion ($\alpha$) were observed at 12 hours (D).

Briefly, hCMVEC cells were seeded into ECIS 96W20IDF plates and cultured in complete M199-hCMVEC supplement serum containing media for 48 hours to allow a stable barrier to form. Cells were then washed thrice with AIM-V-hCMVEC supplement serum-free media and cultured in this media for 16 hours. Media was removed and replaced with respective treatment in AIM-V-hCMVEC supplement serum-free media. Cells were cultured for 24 hours. Barrier properties were monitored in real-time throughout the experiment using ECIS. Ribbon plots represent the mean ± SEM of data from three independent experiments, each consisting of three technical replicates (A-D). Data was normalised 2 hours before the addition of treatment using vascr.

**hCMVEC maintenance:** Human cerebral microvascular endothelial cells (hCMVECs) from Applied Biological Materials Inc (T0259) were cultured in T75 Nunc flasks (Life Technologies, 156499) with M199 medium (Life Technologies, 11150-059) containing 10% FBS (Moorgate), 1μg/mL hydrocortisone (Sigma-Aldrich, H0888), 3ng/mL hFGF (Peprotech, AF-100-18B), 1ng/mL hEGF (Peprotech, AF-100-15), 10μg/mL heparin (Sigma-Aldrich, H3393-50KU), 2 mM GlutaMAX (Invitrogen, 35050061) and 80μM dubutyryl-cAMP (Sigma-Aldrich, D0627) (referred to as M199-hCMVEC supplement serum containing media) at 37°C, with 5% $CO_2$ and 100% humidity. For hCMVEC maintenance and experiments culture vessels were coated with 1μg/cm² collagen I (Invitrogen, C3867) dissolved in 0.02M acetic acid (Merck, 1.00063.2500) for 1 h at room temperature before being washed 3 times with sterile Type I water. Flasks were stored at 4°C for up to 4 weeks before use. Experiments used hCMVEC cells between passages 11 to 16.

Supplementary references

3. Freeman R, Niego B, R. Croucher D, Pedersen LO, Medcalf RL. t-PA, but not desmoteplase, induces plasmin-dependent opening of a blood-brain barrier model under normoxic and ischaemic conditions. *Brain Research* 2014; **1565**: 63–73.
4. Niego B, Lee N, Larsson P, De Silva TM, Au AE-L, McCutcheon F, Medcalf RL. Selective inhibition of brain endothelial Rho-kinase-2 provides optimal protection of an in vitro blood-brain barrier from tissue-type plasminogen activator and plasmin. Wang X, editor. *PLoS ONE* 2017; **12**: e0177332.
5. Niego B, Freeman R, Puschmann TB, Turnley AM, Medcalf RL. t-PA–specific modulation of a human blood-brain barrier model involves plasmin-mediated activation of the Rho kinase pathway in astrocytes. *Blood* 2012; **119**: 4752–61.
