## Supplementary Figure 6 for "Plasmin, the product of tissue plasminogen activator (tPA) treatment for ischemic stroke, impairs human brain endothelial barrier integrity"

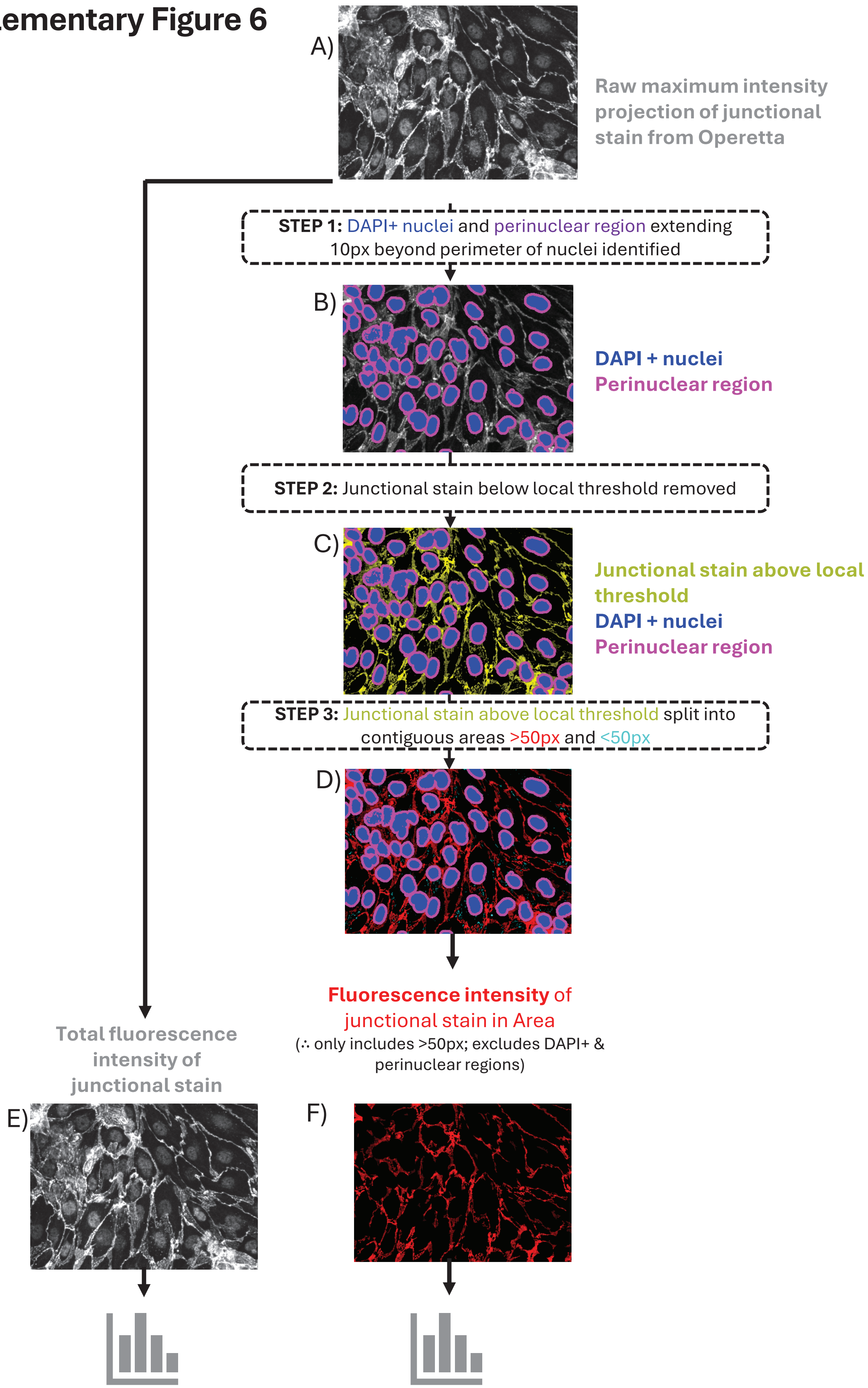

**Supplementary Figure 6. Flow diagram illustrating how immunocytochemistry (ICC) images visualising junctional molecules were processed using vjunctr for quantitative analysis.**

To enable automated quantification of junctional protein expression in large immunocytochemistry confocal image datasets, we developed vjunctr, a custom image-processing workflow. vjunctr improves ICC analysis by enabling objective and reproducible quantification of junctional proteins. It achieves this by excluding nuclear and perinuclear signal, applying adaptive local thresholding to correct for spatial staining variability, and distinguishing continuous junctional networks from punctate cytoplasmic signal. This approach reduces background bias and allows robust, automated comparison of junctional protein expression across large image-datasets.

**Workflow:** hCMEC/D3 cell monolayers were probed with an antibody detecting β-catenin to demonstrate how vjuntcr works.
Raw data consisted of a maximum intensity projection generated from a stack of confocal images captured using an Operetta CLS (A).
**Step 1:** DAPI stain was used to define the nucleus and identify the perinuclear region. A threshold was applied to the DAPI channel to robustly detect nuclei. The perinuclear region was identified by taking the nuclear areas and expanding them by 10 pixels in all directions (B). As nuclei do not overlap with junctional areas in cell monolayers, the nuclear and perinuclear regions were excluded from subsequent analysis of the junctions.
**Step 2:** Identifying antibody staining with intensities above the background (C). Due to the uneven nature of the staining, applying a global threshold across the entire image was unable to reliably discriminate between the stained areas and background (data not shown). Therefore, local thresholding was applied to the image. Local thresholding calculates the difference between each pixel and the average in the local area, thereby correcting for variations in particular areas of the image. To refine this analysis, both the size of the local area and the threshold can be adjusted. For example, selecting a smaller local area will detect smaller objects, whilst a larger area is better suited to detecting larger objects or correcting for gradual changes across the image area. Furthermore, lowering the threshold between each pixel and the local area allows for detection of weaker staining, whilst raising this threshold can compensate for higher background. However, given as the staining strength and background are determined by the immunocytochemical protocol used, rather than the experimental setup or treatments applied, the threshold values can be empirically determined on a small number of representative images, and then automatically applied to large numbers of images. Adjusting these parameters detects both junctional-associated staining and any staining within the cytoplasm (C).
**Step 3**: This step breaks down antibody staining into two groups: i) uniform, uninterrupted networks of antibody staining, and ii) discontinuous, punctate antibody staining (D). These were categorised based upon the area of contiguous staining i.e. >50 pixels or <50 pixels respectively. The contiguous staining >50px was mostly consistent with a junctional pattern, whilst the contiguous <50px punctate staining was predominantly located in the cytoplasm. The >50 px contiguous areas which appeared to represent the cell-to-cell junctional region, were classified as the 'Junctional Area' for subsequent analysis.
To ensure vjuncr processing had identified distinct areas correctly, images were generated at each step for verification. The outputs from the analysis can then be converted into numerical values by vjunctr. The sum of the fluorescence intensities of all pixels within the raw maximum intensity projection represents the total expression of the molecule detected (E). Whilst the fluorescence intensity in the 'Junctional Area' (F) indicates the level of expression at the cell-to-cell junctions. Relative fluorescence measurements can then be automatically compared across large image sets to quantify treatment-dependent changes in junctional protein localisation and expression.
